# Human white matter tracts aligned with canonical functional brain networks

**DOI:** 10.64898/2026.05.20.726546

**Authors:** Kimberly L. Ray, Sohmee Kim, Karen Herrera, Suna Guo, Daniela Reyes, Julio A. Pereza, Anna Bostoganashvili, Gabriele Amorosino, Angela R. Laird, Franco Pestilli

**Author notes:** **Correspondence:** Franco Pestilli.

## Abstract

White matter tracts form the structural backbone of large-scale brain networks, yet their relation to functional organization remains poorly defined. Although major pathways are well characterized anatomically, their relationship to distributed cognitive systems has not been systematically established. Here, we constructed a population-level white matter tract termination atlas and projected tract endpoints onto the cortical surface to characterize their spatial organization. We integrated this atlas with large-scale meta-analytic decoding to derive functional profiles for individual tracts. Functional decoding revealed that white matter tracts exhibited distinct and biologically interpretable cognitive signatures. Hierarchical clustering of these profiles further showed that tracts organize into coherent ensembles defined by shared functional associations. These tract ensembles recapitulated canonical intrinsic brain networks across multiple cortical atlases, including a notable ensemble that demonstrated alignment with the default mode, salience, and frontoparietal control networks, corresponding to the core architecture of the triple-network model of cognitive control. This finding identified a candidate structural backbone linking distributed functional systems implicated across neuropsychiatric conditions. Together, these results demonstrate that white matter architecture is organized according to large-scale functional principles and establish a tract-to-network framework for linking structural connectivity to cognition.

## Introduction

White matter is fundamental to human brain function, providing the long-range structural connectivity that enables communication between distributed cortical and subcortical systems^1,2^. In humans, the expansion and organization of white matter are closely linked to the emergence of higher-order cognition. Advances in diffusion-weighted magnetic resonance imaging (dMRI) and tractography have enabled in vivo reconstruction of major white matter tracts (WMTs), producing increasingly detailed anatomical atlases of white matter organization across individuals^3^.

These studies have established a canonical taxonomy of WMTs based on anatomical trajectory, including association, commissural, and projection pathways^4, 5^. While this anatomical framework has been essential for characterizing the structural architecture of the brain, it does not directly explain how individual tracts contribute to large-scale cognitive systems^6–11^. In parallel, functional neuroimaging has delineated reproducible large-scale cortical brain networks that support perception, action, and higher-order cognition^12,13^. These intrinsic networks have become central to contemporary models of brain function, yet the structural architecture that supports their organization remains incompletely understood.

A central unresolved question is whether large-scale functional systems are supported by individual white matter tracts, coordinated ensembles of pathways, or distributed patterns of connectivity spanning multiple tracts^14,15^. Addressing this question has been challenging for several reasons. Many tracts terminate in broad and overlapping cortical territories, complicating the assignment of discrete functional roles to individual pathways^16^. In addition, substantial inter-individual variability in tract anatomy limits reproducibility and interpretability^17,18^. Existing approaches to structure–function coupling have therefore focused primarily on regional connectivity or whole-brain network models, without directly linking functional organization to anatomically defined tracts.

Recent advances in neuroinformatics provide new opportunities to address this gap. Large-scale meta-analytic frameworks integrate data across thousands of neuroimaging studies to generate probabilistic mappings between brain regions and cognitive processes^12,19,20^. Methods based on machine learning and text mining enable quantitative functional decoding of spatial brain patterns, offering a scalable approach for linking brain organization to cognition^12,21–24^. However, these approaches have been applied primarily to cortical regions, and their potential for characterizing white matter organization remains largely unexplored.

At the same time, work on macroscale cortical organization has revealed that brain function is structured along continuous gradients that span sensorimotor to transmodal systems^25^. These gradients reflect hierarchical transitions from externally driven processing to internally oriented cognition and provide a unifying framework for understanding large-scale brain organization. Whether similar principles extend to the organization of white matter—and whether structural connectivity reflects this hierarchical functional architecture—remains unknown.

Here, we developed a tract-to-network framework that linked white matter pathways to large-scale functional systems. We constructed a population-level atlas of white matter tract terminations mapped onto the cortical surface and combined it with meta-analytic functional decoding to derive tract-level functional profiles. We showed that WMTs exhibit distinct functional signatures and organize into coherent ensembles defined by shared functional associations. These tract ensembles aligned with canonical intrinsic brain networks across multiple atlases. Notably, we identified a tract ensemble whose cortical terminations converge on the default mode, salience, and frontoparietal control networks, revealing a candidate structural substrate for the triple-network model of cognitive control. Together, these findings established a tract-to-network framework that linked white matter architecture to large-scale brain function and provided evidence that white matter organization reflects fundamental principles of human brain organization.

## Results

### White matter tract terminations define a population-level structural framework

We first constructed a population-level white matter tract termination atlas (*WMT*_TA_) by mapping cortical endpoints of major WMTs in 1,062 individuals from the Human Connectome Project^26^ (see **Supplementary Results 1: Supplementary Fig. 1**). For each tract, cortical surface regions of interest were generated to capture the spatial distribution and inter-individual variability of streamline endpoints.

Endpoint distributions varied systematically across tracts. Projection pathways, including the corticospinal tract, and occipital association pathways such as the vertical occipital fasciculus, exhibited spatially consistent and focal termination patterns across individuals. In contrast, long-range association pathways, including the arcuate fasciculus, showed broader and more variable cortical termination zones, particularly within temporal and frontal regions. Smaller tracts displayed increased variability, consistent with prior reports of anatomical heterogeneity (**Supplementary Fig. 1**)^17,18^. Neighboring tracts frequently exhibited overlapping cortical projections, particularly within the occipital cortex, where endpoints of the vertical occipital fasciculus and inferior longitudinal fasciculus converge (**Supplementary Fig. 1**). These overlapping termination zones highlight the challenge of assigning discrete functional roles to individual tracts based solely on anatomical trajectory.

This *WMT*_TA_ provides a surface-based representation of white matter architecture that enables direct alignment with cortical functional organization, establishing a foundation for systematic mapping of tract-level function. The atlas is accessible on brainlife.io (doi: add when published).

### White matter tracts exhibit distinct functional profiles

We next asked whether cortical termination patterns encode tract-level functional specialization. To address this, we integrated the *WMT*_TA_ with large-scale meta-analytic decoding using a Latent Dirichlet Allocation (LDA) framework^20^ trained on thousands of neuroimaging studies^22^. This approach enabled quantitative association of tract endpoints with functional topics. Functional decoding revealed distinct, biologically interpretable profiles across tracts, demonstrating that cortical termination patterns contain sufficient information to infer functional specialization. These profiles emerged without imposing anatomical priors, providing a data-driven mapping between structural pathways and cognitive functions.

To validate this framework, we examined the cortical endpoints of the arcuate fasciculus, a tract with well-established functional organization, and projected them onto surface space (**Fig. 1a**). Bilateral decoding revealed spatially differentiated functional associations corresponding to its cortical terminations, with frontal, superior endpoints associated with language and working memory processes (**Fig. 1b** green) and temporal, inferior endpoints associated with speech-related functions (**Fig. 1b** blue). Hemispheric differences further supported the validity of the approach: the left arcuate showed dominant associations with language-related functions, whereas the right arcuate was more strongly associated with cognitive control and affective processing (**Fig. 1c**). These results recapitulate known functional lateralization patterns^27,28^, indicating that meta-analytic decoding of tract endpoints efficiently captures established neurobiological organization.

**Figure 1.**
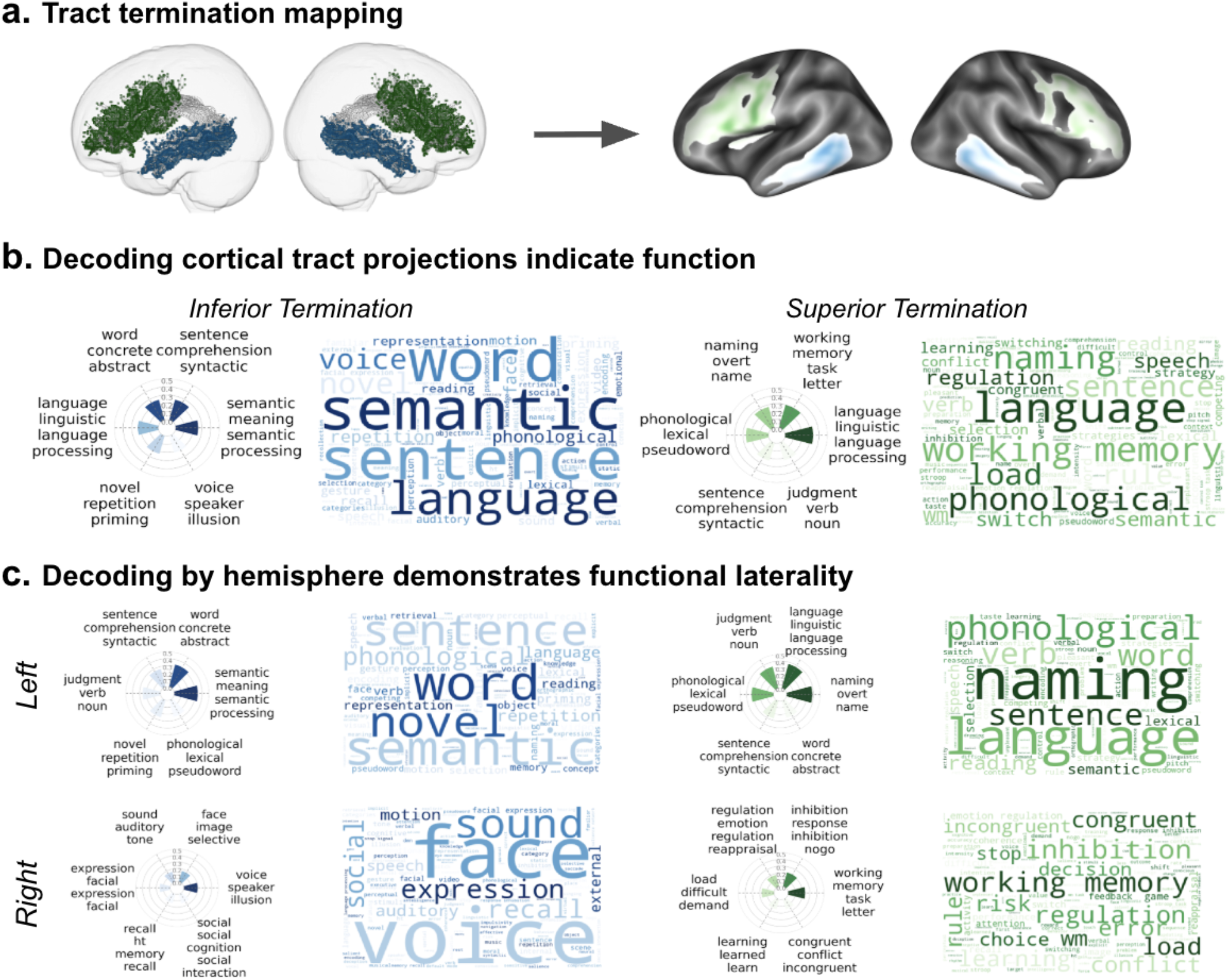
Functions associated with the Arcuate Fasciculus tract terminations: an exemplar framework for functional decoding of white matter tracts. **a**. Cortical termination map and surface cortical visualization of the arcuate fasciculus, highlighting inferior (blue) and superior (green) projection zones of the white matter tract. **b**. Bilateral functional decoding of the arcuate reveals dominant associations with language-related processes. **c**. Hemisphere-specific decoding demonstrates lateralization, with left-hemisphere terminations preferentially associated with language functions and right-hemisphere terminations with socio-emotional processing. Radar plots depict the six highest correlated functional topics for each termination point, and word clouds illustrate the corresponding frequency of terms associated with each termination point. Colors correspond to inferior (blue) and superior (green) projections.

Across all tracts, functional decoding revealed a diverse repertoire of perceptual, emotional and cognitive associations, establishing a comprehensive mapping between white matter architecture and cortical function (**Supplementary Results 2)**. For example, the vertical occipital fasciculus (VOF; **Supplementary Fig. 2**) engaged the dorsal and ventral cortical visual streams with its decoding strongly related to visual processing terms, consistent with its proposed role in integrating spatial and object-based representations^29–31^. Furthermore, the Inferior longitudinal fasciculus (ILF; **Supplementary Fig. 3**), with posterior terminations in the occipital lobe and anterior terminations in the temporal pole^32^, exhibited strong associations with visual–semantic processes, particularly motion and object recognition, in line with its role in linking perceptual inputs to higher-order semantic representations^33–35^. Finally, the corticospinal tract (CST; **Supplementary Fig. 4**) showed highly specific associations with voluntary motor function, including motor planning, imagery, and execution, aligning with its role in motor control^36,37, 38–40^. Together, these findings demonstrate that tract-specific termination profiles give rise to reproducible and functionally coherent signatures, enabling reliable mapping of white matter architecture onto distributed cognitive systems.

### White matter tracts organize into functional ensembles

We next asked whether white matter tracts exhibit higher-order organization based on shared functional profiles. Hierarchical clustering of tract-level functional topic associations revealed a structured and robust organization of white matter. A five-cluster solution was supported by cluster validity metrics, including a high cophenetic correlation coefficient (**Fig 2;** CCC = 0.7362) and favorable separation and compactness measures, indicating that the clustering captures meaningful relationships among tracts (Dunn index; DI = 0.2972; see **Supplementary Results 3, Supplementary Fig. 5**).

**Figure 2.**
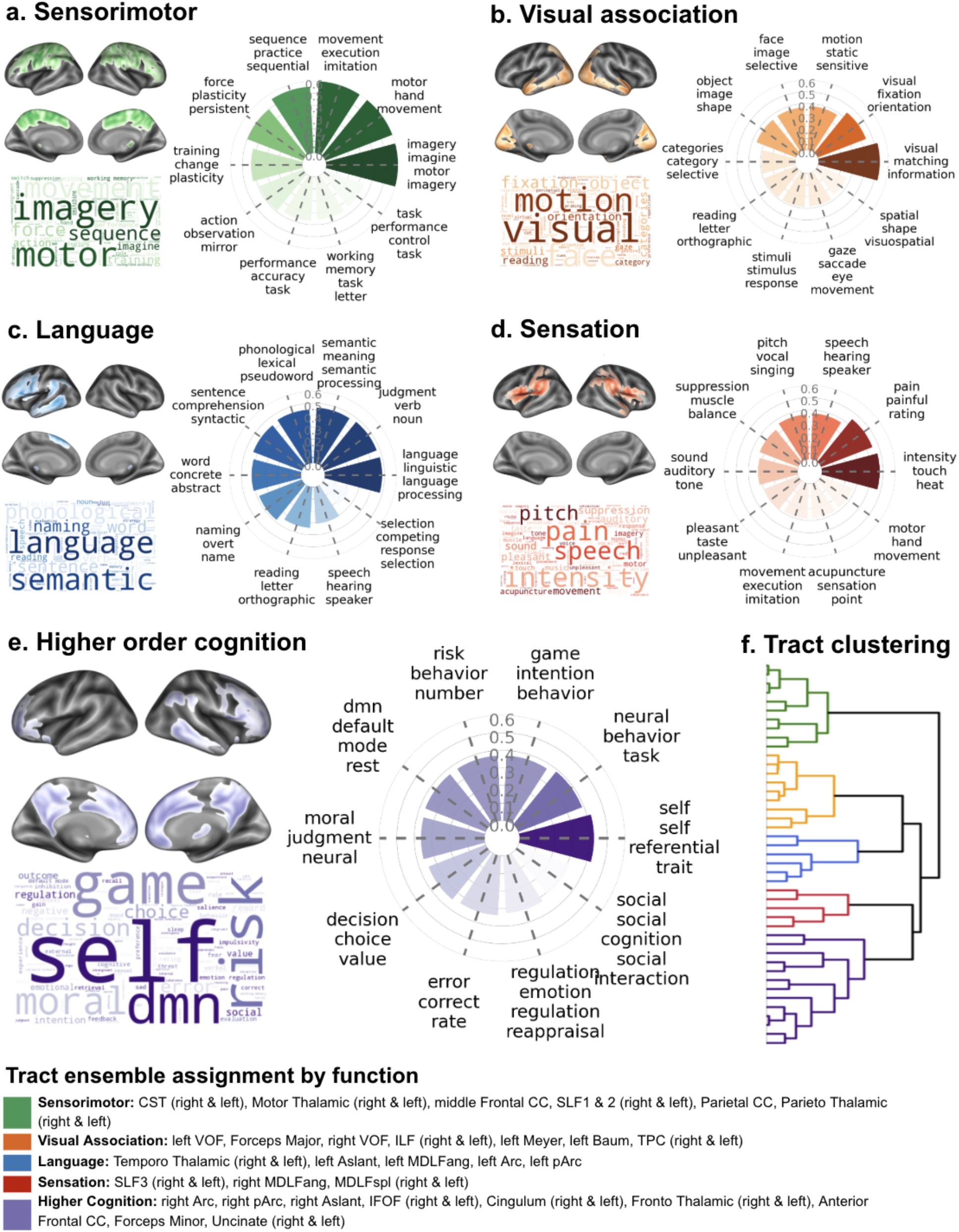
Functional decoding of white matter tract (WMT) endpoint clusters. Functional profiles of the five WMT endpoint clusters identified by hierarchical clustering. For each cluster (**a–e**), aggregated WMT endpoint ROIs were projected onto the cortex. Radar plots display the relative strength of the 10 functional topics most strongly associated with each cluster. Word clouds summarize the dominant functional terms derived from each cluster, providing an intuitive overview of cluster-level functional themes. (**f**) Dendrogram from which the cluster assignments were derived. The list of WMTs assigned to each cluster is listed and color-coded according to cluster membership.

These results indicate that white matter tracts group according to shared functional associations, forming coherent tract ensembles. At the highest level of branching on the dendrogram (**Fig. 2f, Supplementary Fig. 6**), we observed a split between clusters primarily associated with externally oriented, lower-order mental functions (**Fig. 2a**) and clusters associated with more complex, higher-order cognitive functions. A secondary division further distinguished clusters involved in exogenous perception and early information processing (**Fig. 2b and Fig. 2c**) from those supporting endogenous, internally oriented, and goal-directed functions (**Fig. 2d and Fig. 2e**). At finer scales, this organization delineated a progression from motor execution and perceptual processing to increasingly integrative and internally directed cognitive processes. This hierarchical structure parallels known gradients of cortical organization, indicating that white matter architecture is organized along a functional axis that spans sensorimotor to transmodal systems (**Supplementary Results 3** provides detailed descriptions of each cluster).

### Tracts ensembles align with canonical brain networks

Finally, we set to determine whether these novel tract ensembles mapped onto established large-scale cortical brain systems, to do so, we quantified their *spatial alignment* with canonical functional networks across multiple atlases^13,41–47^ using permutation-based spin tests (**Supplementary Results 4; Supplementary Fig. 12**)^48^. Each tract ensemble identified during clustering exhibited significant correspondence with distinct functional networks (spin-test 1,000 permutations for each reference atlas^48^, p<0.05, **Supplementary Table 1-6**). The tract ensemble associated with sensorimotor processing aligned with motor and somatomotor networks (**Supplementary Fig. 13**), while the ensemble associated with visual processing corresponded to visual and dorsal attention networks (**Supplementary Fig. 14**). The language association ensemble mapped onto distributed language and control-related systems (**Supplementary Fig. 15**), and the sensory-interoceptive ensemble aligned with salience and cingulo-opercular networks (**Supplementary Fig. 16**). Across clustering ensembles (**Fig. 2f**), correspondence between white matter tract ensembles and functional networks was consistent across atlases (i.e., the same ensembles typically aligned with similar functional networks), indicating robust structure-function relationships.

Notably, the ensemble of tracts referred to as “higher order cognition” (**Fig. 2e**) aligned with the default mode network (DMN), salience network (SN), and frontoparietal control network (FPN; **Supplementary Fig. 17**). This ensemble comprised thirteen long-range association and projection pathways, including bilateral IFOF, bilateral cingulum, bilateral fronto-thalamic tract, bilateral uncinate fasciculus, the anterior frontal CC, forceps minor, right Aslant, right Arcuate, and right posterior Arcuate, collectively linking the medial prefrontal, posterior cingulate, anterior insular, lateral prefrontal, and medial temporal regions^49^. Together, these three functional networks (DMN, SN, and FPN), comprise the widely studied “triple-network” model of higher-order cognition^49^, in which the SN is proposed to coordinate switching between internally oriented DMN states and externally directed executive processing mediated by the FPN^50–53^. Although the triple-network model has largely been characterized using functional neuroimaging, our findings suggest that interactions among these systems are scaffolded by a shared ensemble-level white matter architecture. Rather than mapping onto isolated tracts, the DMN, SN, and FPN converged onto overlapping sets of long-range structural pathways, suggesting that coordinated tract ensembles may provide an anatomical substrate for dynamic network switching and large-scale cognitive integration. Given the central role of triple-network dysfunction across neuropsychiatric and neurodegenerative disorders, these findings provide a systems-level anatomical framework linking white matter organization to adaptive cognition and its disruption in disease.

## Discussion

A central question in neuroscience is how large-scale functional networks emerge and how white-matter pathways contribute to their organization. Individual networks may depend on multiple tracts, and individual tracts may contribute to multiple functional systems. Here, we establish a tract-to-network framework that links white-matter architecture to the large-scale functional organization of the human brain. By integrating a population-level atlas of tract terminations with meta-analytic functional decoding and hierarchical clustering, we show that white-matter tracts form coherent ensembles with shared functional signatures. These ensembles recapitulate canonical intrinsic brain networks, identifying a tract-level substrate for large-scale cognitive systems.

A primary advance of this work is the demonstration that cortical termination patterns encode sufficient information to infer tract-level functional specialization. Previous studies of structure–function coupling have primarily examined regional connectivity or whole-brain networks^54^, showing that structural connectivity constrains functional interactions. Here, we extend this framework by linking functional profiles directly to anatomically defined tracts. This tract-centric perspective reveals that white matter tracts are not isolated conduits but components of coordinated ensembles that collectively support distributed cognitive processes. The hierarchical organization of these tract ensembles provides further insight into the brain’s macroscale architecture. The progression from sensorimotor and perceptual systems to higher-order, internally oriented cognitive functions parallels established gradients of cortical organization. Prior work has shown that cortical function is organized along continuous axes spanning from externally driven to transmodal processing^25,55^. One implication of these findings is that functional specialization may emerge less from the trajectory of individual white matter pathways than from the coordinated topology of their cortical termination fields. Tracts with distinct anatomical courses frequently converged onto overlapping cortical systems, suggesting that endpoint organization may represent a critical architectural principle linking structural connectivity to large-scale functional dynamics. Prior work has further demonstrated that adaptive cognition depends on dynamic transitions between segregated and integrated large-scale network states during cognitive control and memory processing^56^. In this framework, tract ensembles may support flexible cognition by enabling partially overlapping structural architectures capable of balancing network segregation with rapid integrative processing across distributed cortical territories. This suggests that white matter architecture may contribute to the emergence and stability of macroscale functional gradients^57,58^.

The alignment between white matter tracts and canonical functional networks provides strong evidence that white matter organization supports established large-scale systems. In particular, the identification of a tract ensemble converging on the DMN, SN, and FPN reveals a candidate structural substrate for the triple-network model of cognitive control^49^. These networks are central to models of adaptive cognition, supporting internally directed thought, salience detection, and goal-directed behavior. Our findings suggest that their coordinated function may be facilitated by shared underlying white matter architecture linking their core cortical regions. This structural convergence has important implications for understanding brain dysfunction. Dysregulation of the triple-network system has been independently implicated across neuropsychiatric conditions^59–63^, providing convergent support for the biological relevance of this ensemble-level organization. Consistent with this framework, prior diffusion MRI studies and large-scale meta-analyses have repeatedly reported abnormalities in many of the same white matter pathways comprising the higher-order cognition ensemble, including the cingulum, uncinate fasciculus, inferior fronto-occipital fasciculus (IFOF), arcuate fasciculus, and corpus callosum, across schizophrenia, bipolar disorder, depression, and psychosis-spectrum disorders^64–70^. For example, major depressive disorder has been associated with abnormal salience network organization^59^ and altered interactions with both the DMN and FPN, while mood disorders more broadly exhibit disrupted DMN–FPN coupling linked to rumination and impaired cognitive control^60,71^. Similarly, prodromal psychosis and schizophrenia has been associated with impaired coordination among DMN, SN, and FPN systems^61,72^, alongside white matter abnormalities involving fronto-limbic and fronto-temporal pathways, including the uncinate fasciculus, cingulum, and corpus callosum^64,69^. The present results suggest that vulnerabilities independently identified using functional imaging and white-matter measurements may emerge from a cohesive root cause. Rather than implicating isolated pathways, this framework highlights distributed tract ensembles whose integrity may influence multiple cognitive domains simultaneously, providing a potential structural substrate for transdiagnostic models of brain disorders.

The white matter tracts atlas introduced here stands apart from previous work on white matter atlases. Existing atlases have primarily focused on volumetric representations of tract trajectories^6,7,9,18^, often without precise characterization of cortical endpoints. By mapping tract terminations onto the cortical surface across over 1,000 subjects, the present atlas enables direct alignment with functional organization and supports systematic investigation of white matter structure to cortical function relationships, standing apart from the established function-structural network relationships^54,73^. The observed variability in cortical endpoint-distributions across tracts and individuals further underscores the importance of population-level approaches for distinguishing stable anatomical features from inter-individual variation^16,17^. Despite these advances, several limitations should be considered. Functional decoding relies on large-scale meta-analytic datasets that may reflect biases in the neuroimaging literature and provide statistical associations rather than causal mappings. In addition, dMRI tractography remains constrained by limitations in spatial resolution and the reconstruction of complex fiber configurations^74,75^, which may affect the precision of endpoint localization^76^. Finally, the clustering solution depends on methodological choices that may vary subtly with alternative approaches. Future work should evaluate the stability and generalizability of these tract ensembles across independent datasets, developmental stages, and clinical populations. Integrating microstructural measures, longitudinal data, and genetic information will be critical for understanding the biological mechanisms underlying this organization^77^. More broadly, extending this framework to multimodal datasets may provide a unified account of how structural connectivity, functional dynamics, and cognitive processes are integrated across the human brain.

In summary, this work demonstrates that white matter architecture is organized into functionally coherent ensembles that align with large-scale brain networks. By linking tract anatomy to cognitive function, this tract-to-network framework provides a foundation for investigating how structural connectivity supports cognition and contributes to variability and vulnerability across the lifespan.

## Online Methods

### Generation of white matter termination atlas (WMT_TA_)

Understanding the cortical termination points of white matter tracts is essential for elucidating the structural organization of the human brain and its relationship to function. While existing white matter atlases provide valuable insights into major fiber pathways, they have yet to provide mappings of where these major tracts interface with the cortex. This gap limits our ability to study structure-function relationships, brain connectivity, and neurological disorders that may disrupt these pathways. By creating an atlas that systematically identifies the cortical endpoints of white matter tracts, researchers can enhance neuroanatomical understanding, potentially improve tractography validation, and facilitate studies linking white matter architecture to function, cognitive, and clinical outcomes.

#### HCP Dataset

Anatomical and diffusion MRI data were obtained from the young adult cohort of the Human Connectome Project (HCP) s1200 release^26^. Minimally preprocessed structural T1-weighted images, FreeSurfer reconstructions, and diffusion-weighted images from 1,062 participants were included in subsequent analyses. Detailed acquisition parameters and preprocessing procedures are described in prior HCP publications^78,79^.

#### MRI Data Analysis

##### Tissue segmentation and white matter tractography

Anatomical and diffusion images from the HCP dataset were processed as part of a previous study^80^ using brainlife.io. A complete list of brainlife.io applications and version numbers used in the processing pipeline is provided in **Supplementary Table 7**. The current work utilized 61 white matter tractograms provided in Hayashi et al (2024) to map the tract termination points to cortical surface. Analysis steps used to create white matter tractograms were based on MRTrix functions and are outlined in detail in Hayashi et al. 2024 (**Supplementary Fig. 1, first column**).

##### Mapping white matter termination points to cortical surface

Tractograms underwent a series of processing steps to generate cortical termination maps (**Supplementary Fig. 1, second column**). This process began with removing outlier WM streamlines from tractograms using functionalities in Vistasoft, executed through brainlife.io (centroid standard deviation = 4, length standard deviation = 4, maximum number of iterations = 5; DOI: 10.25663/brainlife.app.195). This step prepared and cleaned data for accurate cortical projection. Next, white matter streamline endpoints of the cleaned tractograms were mapped onto the gray matter surface to generate white matter termination atlases via the Freesurfer mri_convert functionality, implemented on brainlife.io (Gaussian decay function, decay radius threshold = 1; DOI: 10.25663/brainlife.app.194). This step terminated endpoints extending beyond the cortex and created a unique map/image for each end of all 61 WMTs, resulting in 137 images corresponding to the left/posterior/inferior (LPI) and right/anterior/superior (RAS) endpoints.

The endpoint maps were then registered to a common space. In this process, ROIs (regions of interest) were resliced to align voxel grids with input anatomy. This was implemented on brainlife.io using Freesurfer’s mri_vol2vol function (DOI: 10.25663/brainlife.app.522). Next, all 137 endpoint maps for each subject were warped from native space to standard MNI152 space (1.25mm isotropic voxels) using FSL flirt and fnirt functionalities. The resulting maps were then binarized to create individual ROIs (DOI: 10.25663/brainlife.app670).

Next, volumetric endpoint maps were mapped to surface space using Freesurfer’s mri_vol2surf function (DOI: 10.25663/brainlife.app.759). Operating on the endpoint maps in surface space avoided the potential inaccuracies in template warping and produced a better endpoint atlas. After transforming to surface space, the endpoint maps were smoothed using Connectome Workbench -metric-smoothing function (kernel type = GEO_GAUSS_AREA, kernel size = 3; DOI: 10.25663/brainlife.app768).

Finally, endpoint maps for each WMT were averaged across subjects using the Connectome Workbench’s - surface-average and -metric-math functions. The aggregation of averaged WMT endpoints maps produced a density atlas for each cortical termination of each of the 61 major WMTs. Each map represented, for a single ROI of a WMT, the percentage of subjects whose ROI landed at that surface vertex/location. Sixteen WMTs were excluded because at least one end of these tracts terminated in subcortical regions (e.g., spinothalamic and cerebellar tracts). Lastly, we applied a threshold of 0.15 to exclude low-probability endpoint locations, retaining the top 85% of surface vertices where subjects exhibited tract terminations. Furthermore, visual inspection of the endpoint maps revealed a wide range of anatomical variation across subjects. Substantial anatomical variability for two tracts (right Baum and right Meyer) was observed, which were excluded from subsequent analyses. The endpoint maps from the remaining 43 WMTs underwent subsequent LDA functional decoding.

### LDA-based meta-analytic maps

Functional decoding of white matter tract endpoints was performed using a meta-analytic Latent Dirichlet Allocation (LDA) model trained on the NeuroQuery database^22^ and implemented in the gradec Python library^81^ (**Supplementary Fig. 1, third column**). The LDA model identified 200 semantically coherent neuroscience-related topics derived from thousands of published neuroimaging studies^20^.

Topics categorized as anatomical or clinical were excluded, leaving 104 functional topics for analysis. Each white matter tract endpoint map was correlated with the 104 meta-analytic topic maps using Pearson’s correlation. Functional decoding produced three outputs for each tract endpoint: (1) correlation values across topics, (2) radar plots showing the strongest topic associations, and (3) word-cloud visualizations summarizing dominant functional terms (**Supplementary Fig. 1, fourth column**).

### Hierarchical clustering analysis

Hierarchical clustering analysis (HCA) provided a robust and interpretable framework to reveal hidden functional clusters that might otherwise be obscured by traditional analyses. Using this approach, we grouped data into hierarchically structured clusters based on a similarity metric, generating a dendrogram that visually represented hierarchical relationships. Because HCA did not require a priori specification of cluster numbers, it was particularly well-suited for exploratory analysis of high-dimensional data.

To cluster the 43 WMTs, endpoint maps for each tract (RAS + LPI) were combined into a single image and then submitted to the LDA decoder for functional assessment of each WMT. Using the functional decoding topics from the LDA-decoder, HCA was performed on a correlation matrix of 104 functional topics and 43 major WMTs. The distance between clusters was defined as 1 – r, where r is Pearson’s correlation coefficient of WM endpoint maps and functional topics. The complete linkage function from the scipy package was used to generate compact clusters that minimize intra-cluster distances and maximize inter-cluster distances.

A dendrogram was used to visualize HCA results, stratifying WMTs according to the similarity of their endpoint functional topic associations. To evaluate the robustness of the clustering solution, the cophenetic correlation coefficient was computed. This provided a quantitative measure of how well the hierarchical clusters preserved the original pairwise distances within the dataset. To determine the optimal number of clusters, the Dunn index, an evaluation metric for clustering quality, was computed. Finally, functional decoding, via the *gradec* library, was carried out on each of the resultant clusters from the HCA solution.

### Functional Network Correspondence

We next examined whether the white matter tract ensembles identified through HCA aligned with known canonical large-scale functional networks (e.g., DMN, dorsal attention, FPN). These analyses allowed us to test whether structurally distinct tract groupings recapitulate known patterns of functional integration, consistent with the notion of structure-function coupling^82,83^. If significant correspondence was observed, it would further support that macroscale white matter architecture was organized in a manner that supported and potentially constrained the topology of functional brain networks.

To quantify this correspondence, we employed the Network Correspondence Toolbox (NCT^48^), which provided an efficient permutation-based framework to compute spatial similarity metrics between brain maps (**Supplementary Fig. 12**). Specifically, we calculated Dice coefficients for each white matter tract cluster against a series of widely used functional atlases, with statistical significance assessed via nonparametric spin tests (1,000 for each specific atlas tested) to control for spatial autocorrelation. The following reference atlases were included in our analysis: EG17^13,41^, TY7^42^, AS200Yeo17^42,43^, MGlasser360J12^44,45^, XS268_8^46^, and EG286_12^13,47^. Each comparison yielded a Dice coefficient and associated spin-based p-value, providing a robust quantification of structure-function alignment across multiple functional frameworks.

## Acknowledgements

This research was supported by the following grants: Wellcome (226486/Z/22/Z, Principal Investigator F. Pestilli); NINDS UM1NS132207, BRAIN CONNECTS: Center for Mesoscale Connectomics (Principal Investigator K. Ugurbil); and NINDS U24NS140384, BRAIN CONNECTS: The Axonal Projectome EXchange (APEX) (Principal Investigator F. Pestilli). We thank Amazon Web Services Open Data Sponsorship Program for supporting data storage for brainlife.io

