## Supplementary Material for "Human white matter tracts aligned with canonical functional brain networks"

### Abstract

White matter tracts form the structural backbone of large-scale brain networks, yet their relation to functional organization remains poorly defined. Although major pathways are well characterized anatomically, their relationship to distributed cognitive systems has not been systematically established. Here, we constructed a population-level white matter tract termination atlas and projected tract endpoints onto the cortical surface to characterize their spatial organization. We integrated this atlas with large-scale meta-analytic decoding to derive functional profiles for individual tracts. Functional decoding revealed that white matter tracts exhibited distinct and biologically interpretable cognitive signatures. Hierarchical clustering of these profiles further showed that tracts organize into coherent ensembles defined by shared functional associations. These tract ensembles recapitulated canonical intrinsic brain networks across multiple cortical atlases, including a notable ensemble that demonstrated alignment with the default mode, salience, and frontoparietal control networks, corresponding to the core architecture of the triple-network model of cognitive control. This finding identified a candidate structural backbone linking distributed functional systems implicated across neuropsychiatric conditions. Together, these results demonstrate that white matter architecture is organized according to large-scale functional principles and establish a tract-to-network framework for linking structural connectivity to cognition.

### Supplementary Results 1: White matter termination atlas ($WMT_{TA}$)

We constructed a cortical surface–based white matter tract termination atlas ($WMT_{TA}$) using diffusion MRI data from 1,062 individuals in the Human Connectome Project[1]. Across tracts, we defined 137 surface-based regions of interest (ROIs) corresponding to endpoint distributions.

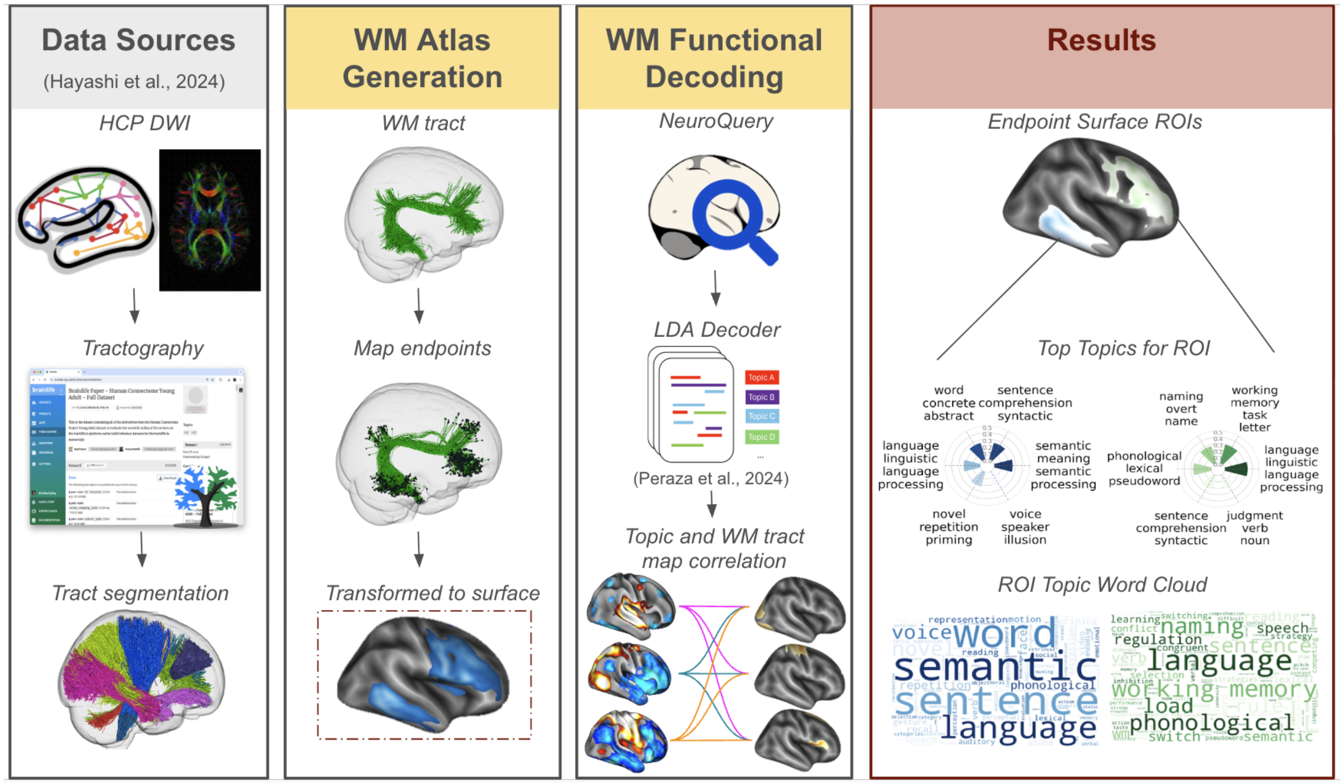


**Supplementary Figure 1. Overview of Analysis Pipeline**. **Data Sources** (Left): Segmented tractograms from 1,062 HCP participants, published in Hayashi et al., 2024, were obtained for analysis. **Methods** (Middle, Left Column) The segmented tractograms were used to map white matter termination points to cortical gray matter regions in individual space and standardized to MNI space. Termination points were averaged across all subjects to create an average density atlas. (Middle, Right Column) Text and brain activation coordinates were extracted from previously published studies using Neuroquery (Dockès et al., 2020). LDA methods outlined in Pereza et al. (2024) were applied to Neuroquery outputs and integrated with the density atlas to produce 104 functional topic-based maps. **Results** (Right) LDA analysis quantified the correlation between each WMT endpoint and the 104 identified functional topics from NeuroQuery.

Variation in the distribution properties of endpoints across the tracts were observed (**Supplementary Fig 2**). Larger WMTs typically produced larger WMT ROIs, similarly, smaller WMTs tended to produce smaller ROIs. However, the density distribution of the endpoint ROIs was not directly associated with WMT volume. For example, major tracts like the CST and vertical occipital fasciculus (VOF) showed tighter distribution (i.e., less variability across participants) of termination locations, while arcuate fasciculus showed wider distribution (i.e. greater variability across participants). Less prominent WMTs such as the right baum (optic radiation) and right meyer also showed greater variability in their endpoint ROIs across subjects.

Further, it was common to observe overlap in the endpoints of neighboring ROIs, which has been demonstrated previously[2,3]. For example, posterior endpoints of VOF and ILF visual tracts demonstrated some overlap in the occipital lobe (see **Supplementary Fig 2**). A brief discussion of the anatomical locations of select exemplar endpoint ROIs (**Supplementary Fig 2**) and their consistency with prior literature is provided below.

We note that the right baum and right Meyer tracts were removed from subsequent functional decoding because of their high variability across participants.

**Arcuate Fasciculus:** Termination points for the arcuate fasciculus were located in the frontal and temporal lobes. The superior endpoints were larger in the left hemisphere than the right, extending to the lateral portions of the precentral, mid frontal, and superior giri, notably, Broca's area (BA 44). Inferior arcuate endpoints were located in posterior areas of the temporal lobe, including Wernicke's area (BA 22). Both endpoint locations are consistent with previous post-mortem tract tracing and DTI studies[4–6].

**Vertical Occipital Fasciculus (VOF):** Both the superior and inferior termination points were located in the occipital lobe. The inferior VOF endpoint was located in the basal occipital area, including V8, V4, the occipital face area, and the fusiform face area. These results are consistent with post-mortem dissection and tractography studies [7–9]. More specifically, the superior VOF endpoint was located in V3 and V7 visual areas. Notably, the VOF endpoint map did not include VI striate cortex.

**Inferior Longitudinal Fasciculus (ILF):** The posterior ILF termination points were located in the occipital lobe and encompassed both striate and extrastriate cortices except the calcarine fissure. The anterior ILF termination points, located closer to the temporal pole (BA 38), extend to the superior temporal gyrus (part of BA 22) and middle temporal gyrus, but not the inferior temporal gyrus. Previous tract tracing and DTI research have demonstrated that the ILF connects the fusiform gyrus within the occipital and temporal lobes [10–12].

**Corticospinal Tract (CST)**: Only the anterior termination points of the corticospinal tract (CST) were considered, as the posterior portion of the CST terminated in the spinal cord, falling outside the scope of our cortical analysis. The anterior termination points were located in the primary motor cortex, superior primary somatosensory cortex, and posterior parietal cortex, aligning with the CST's well-established termination patterns[13,14].

**Optic Radiation (**Baum**)**: Baum's loop[15] is commonly referred to as the posterior/dorsal component of the optic radiation[16]. The anterior endpoint of this tract was located in the lateral geniculate nucleus (LGN), a relay nucleus of the visual pathway. The posterior endpoint was located in the superior occipital cortex. We note that Baum's posterior endpoint in particular demonstrated a high level of subject variability; this can be seen in **Supplementary Figure 2,** where the unthresholded endpoint spreads across the entire occipital lobe, yet when eliminating voxels containing less than fifteen percent of individual participants' WMT endpoints, the tract is substantially smaller, only residing in the left hemisphere. Origins of this tract are not well established[16] and as a likely consequence, there is not a substantial body of work demonstrating its anatomical location. The observed variability across participants in the current study may be an indication of why it is not well studied/described in the neuroscience literature.

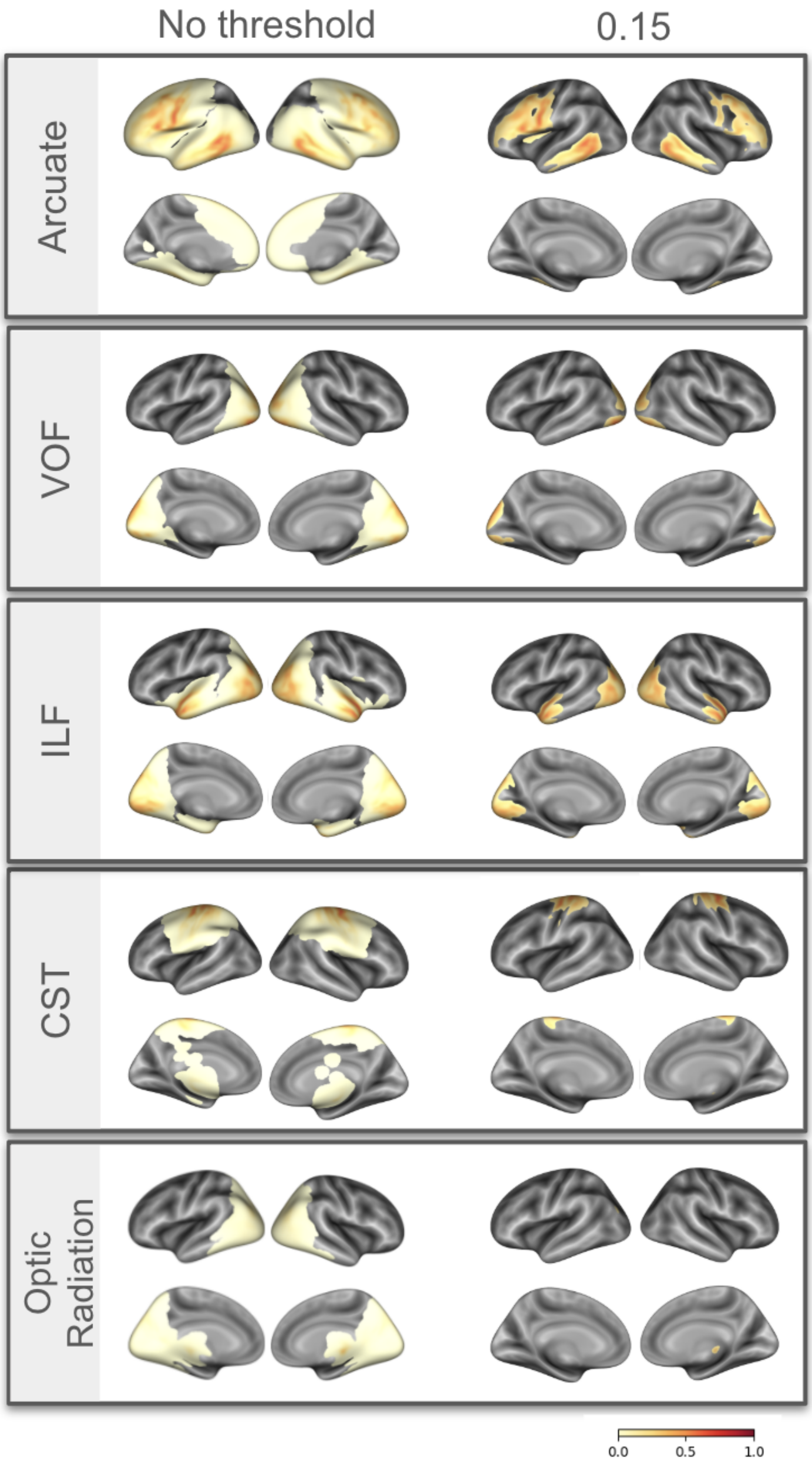


**Supplementary Figure 2:** Exemplar white matter tract endpoint density maps of the arcuate, VOF, ILF, CST, and Baum. Each endpoint density map is shown unthresholded (left), showing the full range of subject variability, and thresholded (right), where voxels containing less than 15% of endpoints are removed.

**Supplementary Results 2: LDA-based Meta-analytic functional decoding of tract endpoints**

We next asked whether cortical termination patterns encode tract-level functional specialization. To link structural connectivity with cognitive function, we applied latent Dirichlet allocation (LDA)–based decoding using NeuroQuery-derived meta-analytic maps. Endpoint ROIs were decoded individually and in bilateral combinations, and functional associations were visualized using word clouds and radar plots summarizing dominant topic correlations.
This approach revealed functionally distinct profiles across tracts and along their anterior–posterior axes (**Fig. 1; Supplementary Figs 3-9)**. Meta-analytic functional decoding of WMT$_{TA}$ ROIs is provided below. For each WMT$_{TA}$ ROI, the word cloud visualized the correlation of individual terms with respect to the ROI, with darker color and larger size of text representing higher correlation values. Complementary to the word clouds, the radar plots depicted the six topics with the strongest correlation values identified during LDA decoding, offering a higher-level thematic summary of functional associations.
The full repertoire of WMT$_{TA}$ endpoint ROIs and their functional decoding was carried out and can be accessed below.

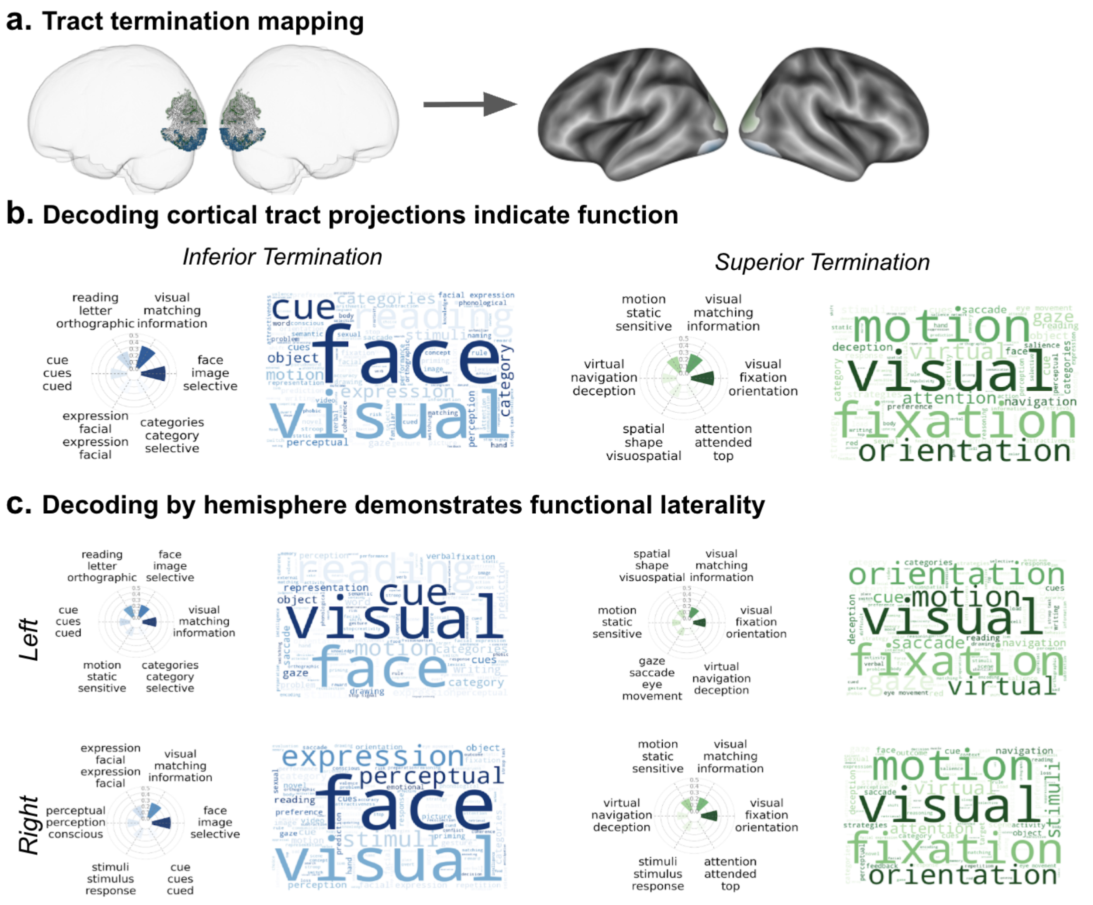


**Supplementary Figure 3.** Functions associated with the termination points of the Vertical Occipital Fasciculus (VOF). **a**. VOF endpoint mapping from the WMT$_{TA}$. **b**. Surface plot showing the anterior and posterior termination points of the white matter tract. **c**. bilateral functional decoding, **d**. Left hemisphere decoding, and **e**. Right hemisphere decoding of the VOF, including a word cloud illustrating the frequency of terms associated with each termination point and a Radar plot depicting the six highest correlated functional topics for each termination point.

**Functional decoding of the Vertical Occipital Fasciculus (VOF) terminations**

First, we examined the cortical endpoints of the VOF (**Supplementary Figure 3**) and its functional decoding. The inferior termination points (blue) of the VOF primarily engaged the ventral visual stream, while the superior termination points (green) were associated with the dorsal visual stream. The word cloud and radar plots reinforced these findings, highlighting that both termination points are strongly related to visual processing. The superior termination points (blue) showed a greater association with spatial awareness and motion guidance, involving topics such as virtual-navigation-deception and gaze-saccade-eye movement. In contrast, the inferior termination points (green) were more strongly connected to the perception and recognition of shapes, objects, and faces, including topics such as face-image-selective and reading-letter-orthographic. No notable differences in functional decoding of right and left VOF tracts were noted. These results align with previous research on the VOF's crucial role in integrating visual information and its involvement in connecting the ventral and dorsal visual streams[7,17,18].

#### Functional decoding of the Inferior Longitudinal Fasciculus (ILF) terminations

Next, we examined the cortical endpoints of the ILF (**Supplementary Figure 4**) and its functional decoding. The inferior longitudinal fasciculus (ILF) demonstrated a stronger functional division between its anterior and posterior termination points. The posterior ILF termination points, located in the occipital lobe, were strongly associated with visual processing, particularly in tasks related to motion detection, gaze/saccade, and object recognition. These associations underscore the ILF's role in visual functions. Previous tract tracing and DTI research have demonstrated that the ILF connects the occipital and temporal lobes[10], facilitating the integration of visual, auditory, and semantic processing[19–21]. Decoding of these regions presents similar functional characterizations of the ILF. More specifically, the anterior ILF termination points showed strong links to auditory processing and some higher-order cognitive functions.

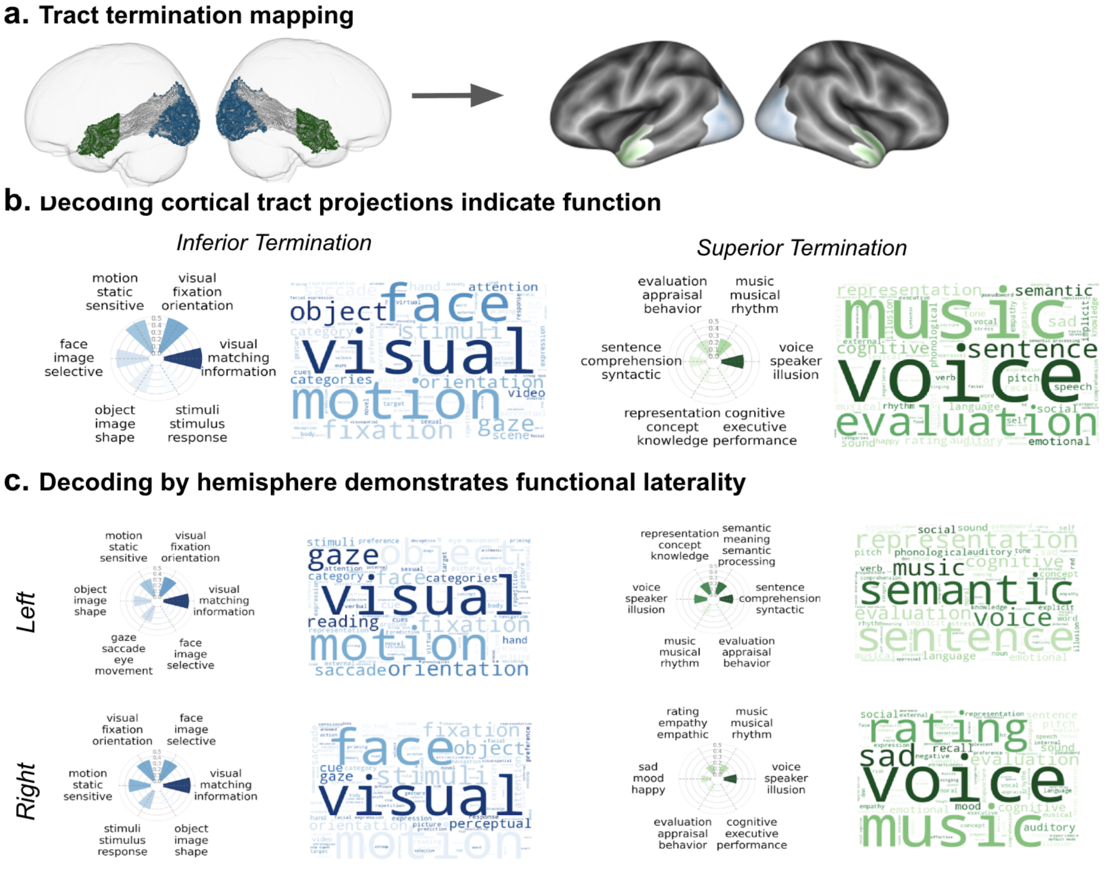


**Supplementary Figure 4. Functions associated with the termination points of the Inferior Longitudinal Fasciculus (ILF). a**. ILF endpoint mapping from the $WMT_{TA}$. **b**. Surface plot showing the anterior and posterior termination points of the white matter tract. C. bilateral functional decoding, d. Left hemisphere decoding, and **e.** Right hemisphere decoding of the ILF, including a word cloud illustrating the frequency of terms associated with each termination point and a Radar plot depicting the six highest correlated functional topics for each termination point.

The word cloud and radar plot (**Supplementary Figure 4**) emphasize the anterior ILF's involvement in auditory functions, such as speech perception, music processing, and semantic understanding. Additionally, these anterior termination points were associated with cognitive tasks like sentence comprehension, concept formation, and semantic processing. These findings suggest that the anterior ILF plays a role in both auditory processing and the integration of sensory information for higher-level cognitive functions, including semantic processing and executive functions.

#### Functional decoding of the Corticospinal Tract (CST) terminations

Finally, we examined the cortical endpoints of the CST (**Supplementary Figure 5**) and its functional decoding. Bilateral functional decoding of the CST highlighted its well-established role in voluntary motor control[22,23]. More specifically, the CST was most strongly related to topics that described voluntary movement, motor planning, imagining, and execution, as well as the body's fine motor control, including hands and limbs. Stroke and motor learning research further support the function of this WMT[24–26]. Functional associations for right and left CST were highly consistent with these findings. The prominence of motor-related terms in the word cloud further supports the CST's role in fine-tuned coordination and skilled movement, while the radar plots provide a structured view of functional domains associated with these termination points.

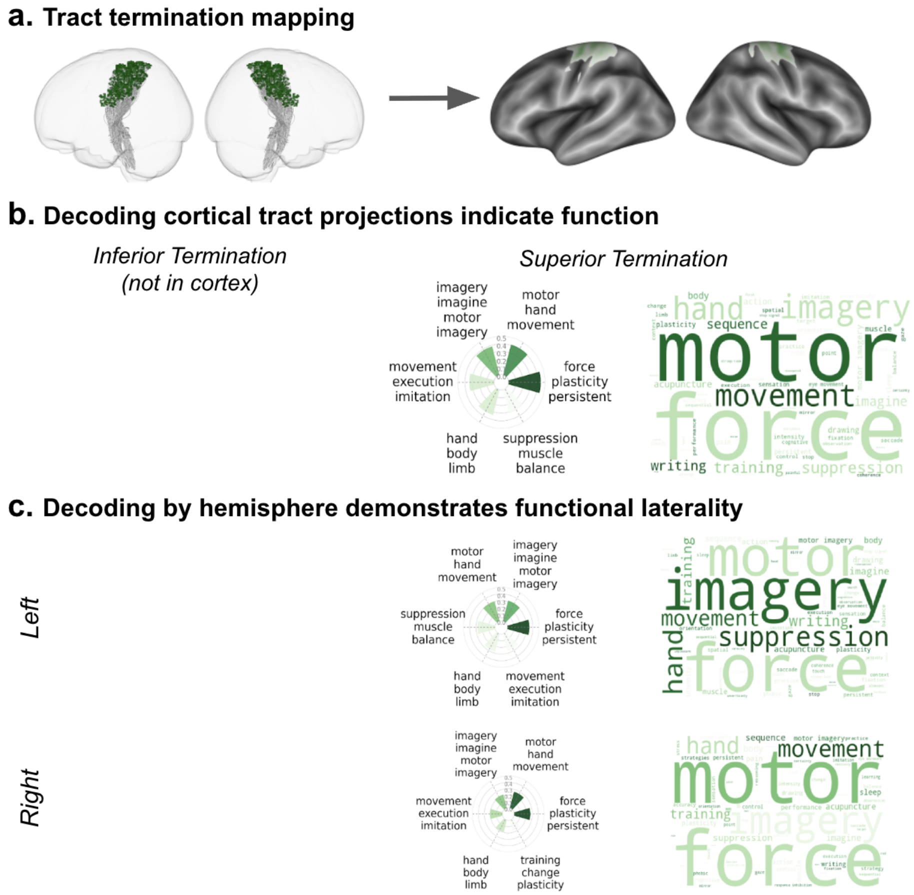


**Supplementary Figure 5.** Functions associated with the cortical termination points of the corticospinal tract (CST). **a**. CST endpoint mapping. **b**. Surface plot of the anterior termination points of the white matter tract, only the anterior termination point is visualized, as the posterior terminations extend into the spinal cord. **c**. bilateral functional decoding of the CST, including a word cloud illustrating the frequency of terms associated with each termination point and a Radar plot depicting the six highest correlated functional topics for each termination point.

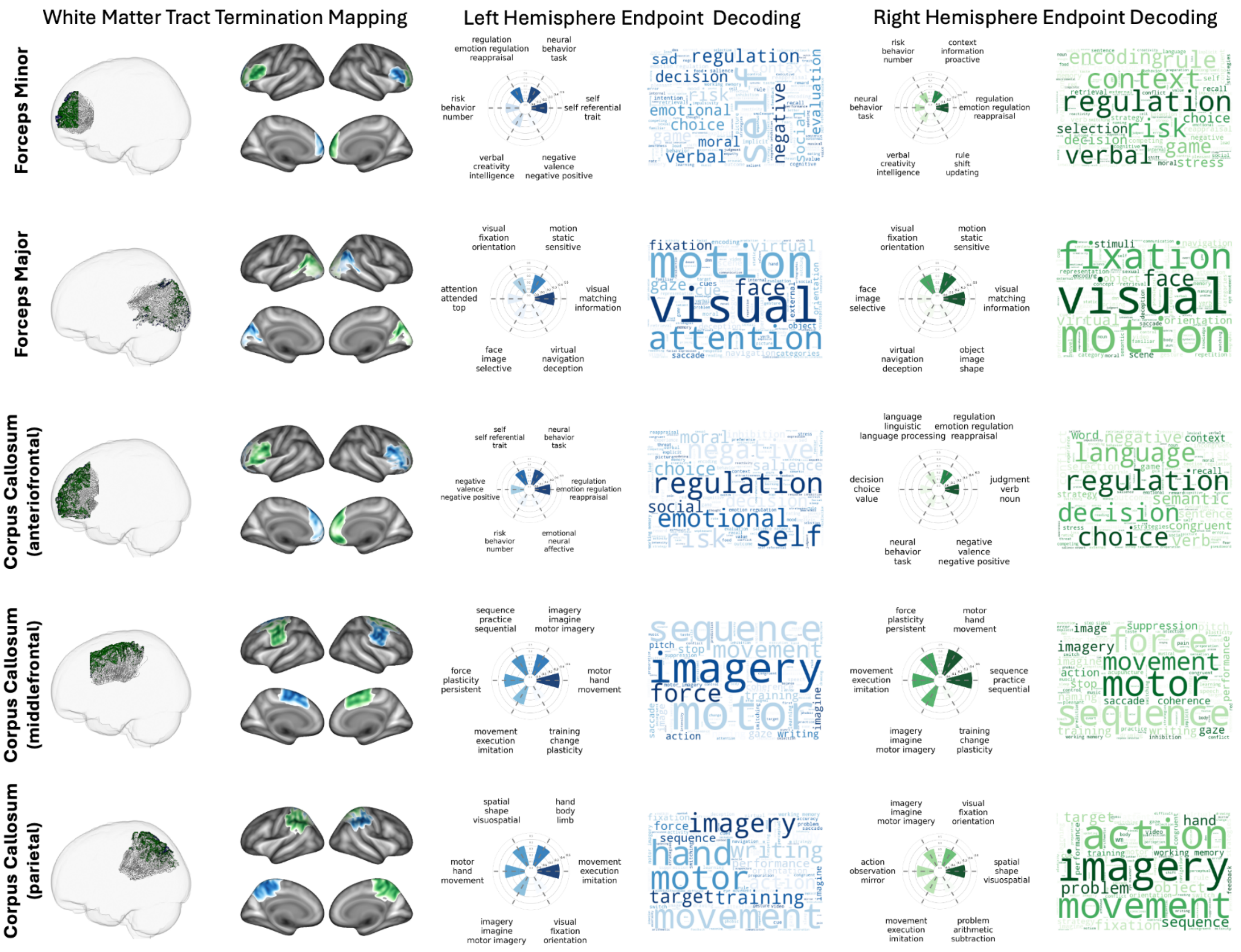


**Supplementary Figure 6. Functional decoding of cortical termination points for midline white matter tracts.** Cortical termination maps and associated functional profiles are shown for the following midline white matter tracts: Forceps Minor, Forceps Major, Anterior Frontal Corpus Callosum, Middle Frontal Corpus Callosum, and Parietal Corpus Callosum. Because these tracts traverse the midline, each tract contains a single cortical termination region within each hemisphere. The left columns display cortical endpoint distributions derived from the White Matter Tract Termination Atlas ($WMT_{TA}$). The right columns present functional decoding results for the left and right hemisphere termination regions, including word clouds illustrating the relative frequency of functional terms associated with each cortical endpoint and radar plots summarizing the six highest-correlated functional topics for each termination region.

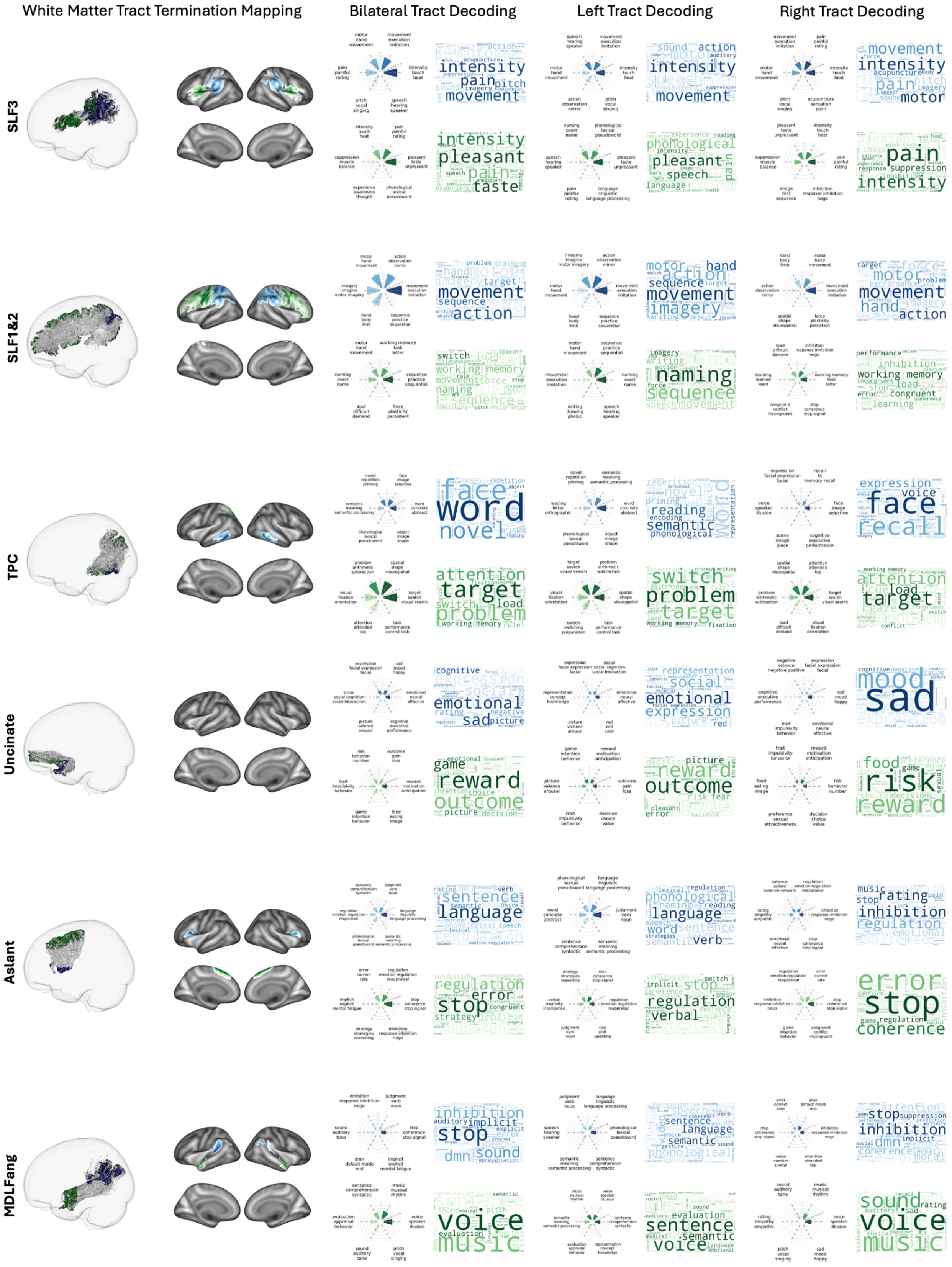
White Matter Tract Termination Mapping
Bilateral Tract Decoding
Left Tract Decoding
Right Tract Decoding
SLF3
SLF1&2
TPC
Uncinate
Aslant
MDLFang

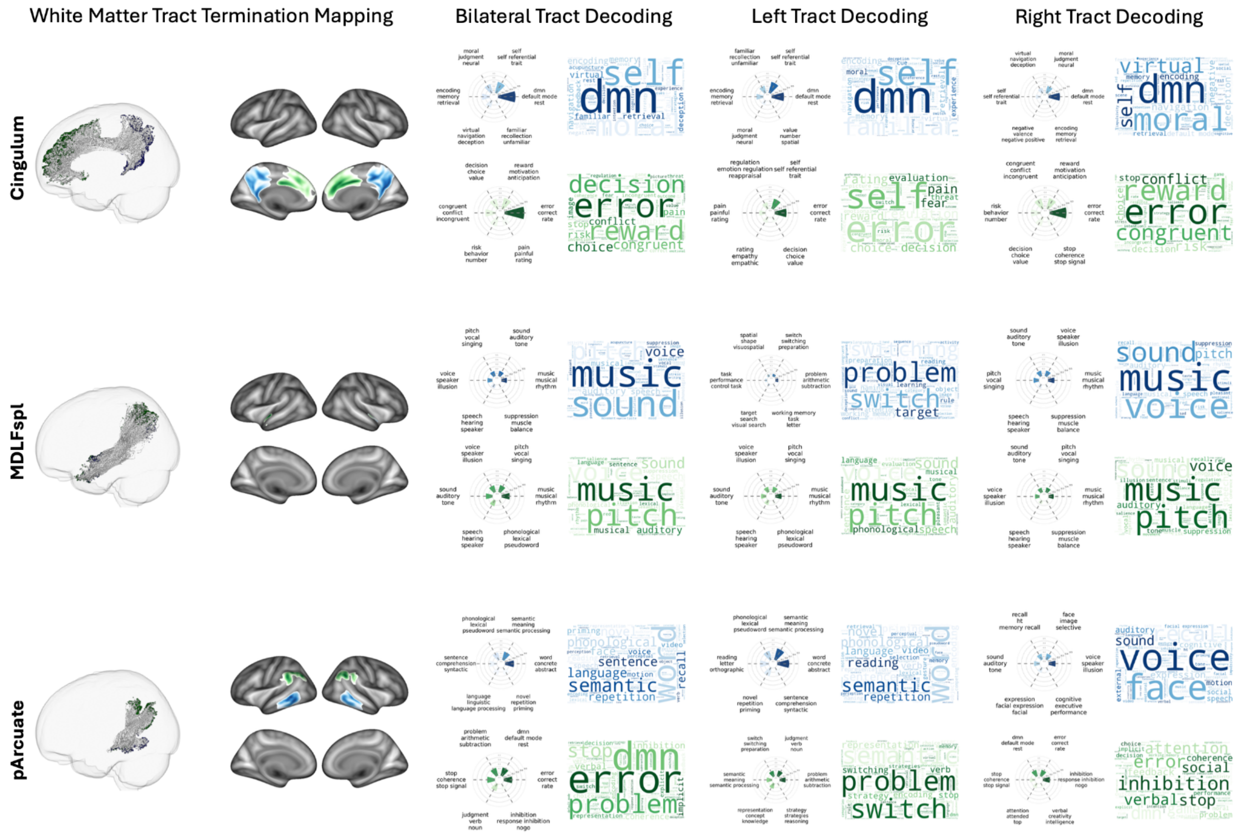


**Supplementary Figure 7. Functional decoding of cortical termination points for association white matter tracts.** Cortical termination maps and associated functional profiles are shown for the following association white matter tracts: MDLFang, posterior Arcuate, SLF I&II, SLF III, Uncinate, Aslant, Cingulum, MDLFspl, and TPC. Functional decoding is presented for the bilateral tract terminations (combined left and right WMT$_{TA}$ ROIs), as well as separately for the left hemisphere and right hemisphere tract termination regions. The left columns display cortical endpoint distributions derived from the WMT$_{TA}$. The right columns present functional decoding results for each termination region, including word clouds illustrating the relative frequency of functional terms associated with each cortical endpoint and radar plots summarizing the six highest-correlated functional topics for each termination region.

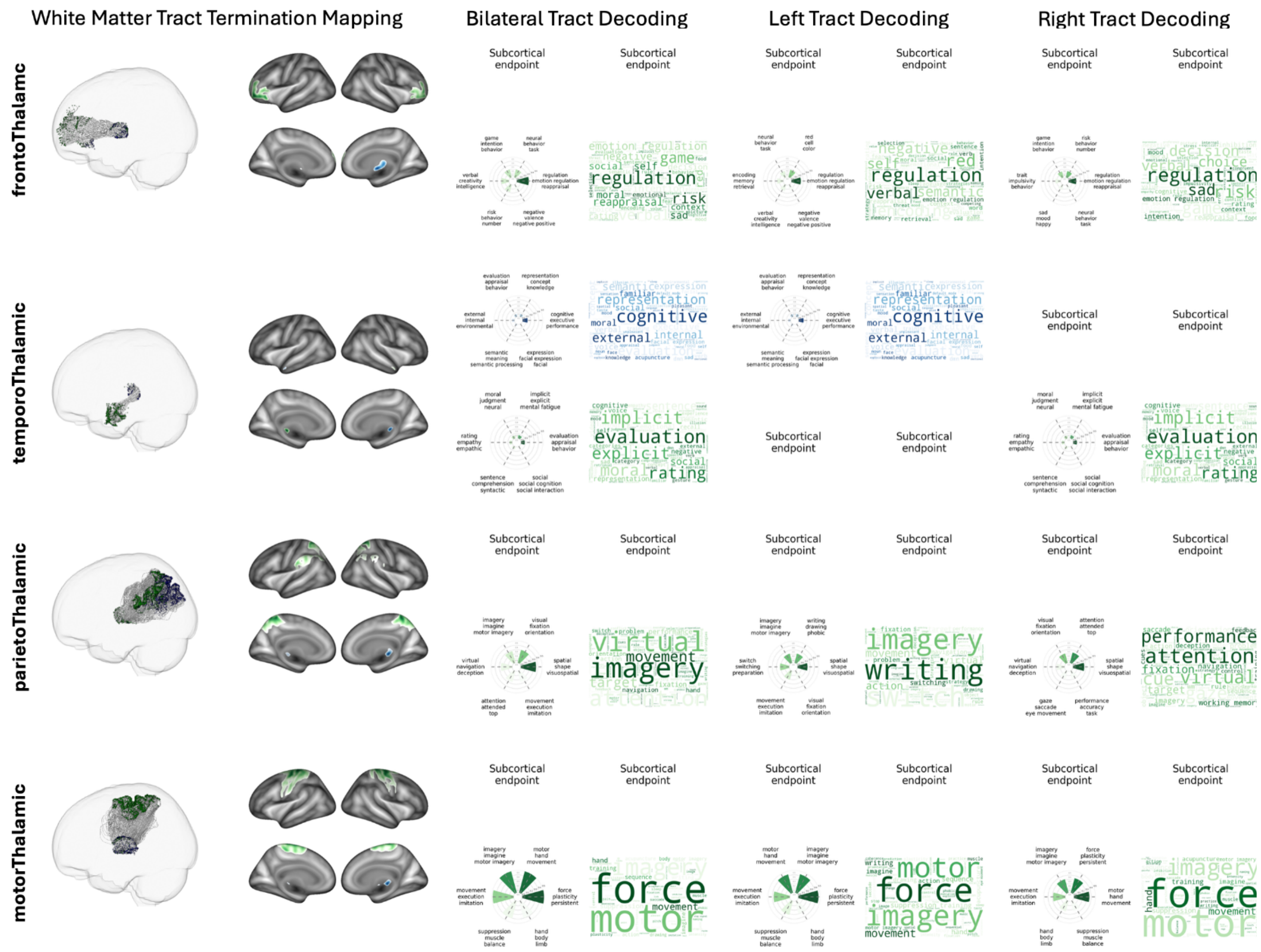


**Supplementary Figure 8. Functional decoding of cortical termination points for thalamic white matter tracts.** Cortical termination maps and associated functional profiles are shown for thalamic white matter tracts. Because these tracts end at the thalamus, each tract contains a single cortical termination region within each hemisphere. The left columns display cortical endpoint distributions derived from the $WMT_{TA}$. The right columns present functional decoding results for each termination region, including word clouds illustrating the relative frequency of functional terms associated with each cortical endpoint and radar plots summarizing the six highest-correlated functional topics for each termination region.

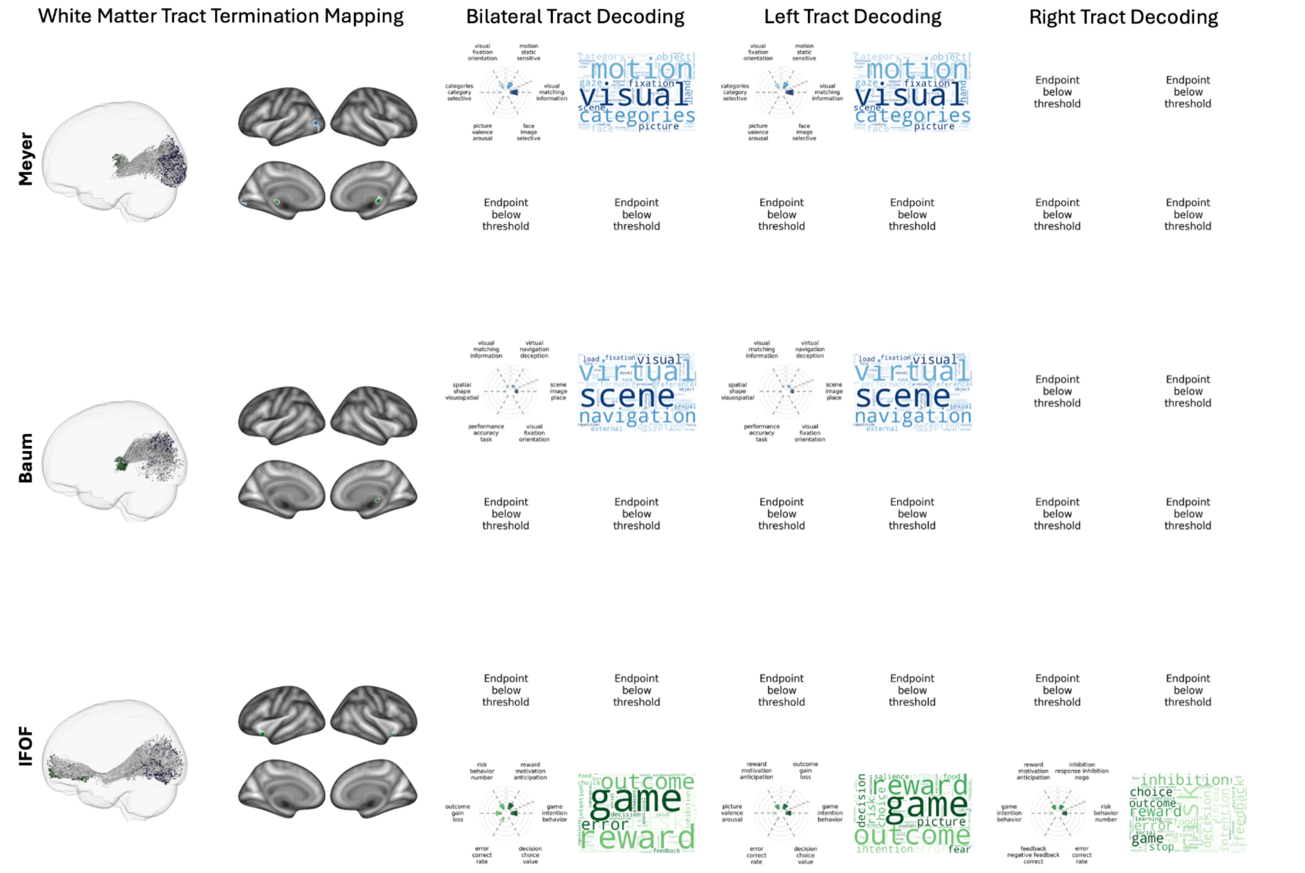


**Supplementary Figure 9. White matter tracts excluded from functional decoding analyses.** Cortical termination maps are shown for the remaining white matter tracts that did not meet thresholding criteria for subsequent functional decoding analyses, including Meyer's loop, posterior optic radiation (Baum), and the inferior fronto-occipital fasciculus (IFOF). Meyer's loop and posterior optic radiation endpoint maps produced $WMT_{TA}$ ROIs that were too spatially limited or sparse to support reliable functional decoding. As a result, these tracts were excluded from downstream meta-analytic functional characterization. The posterior terminations of the IFOF did not meet thresholding and thus only the anterior terminations were included for analysis. The displayed maps illustrate the cortical endpoint distributions derived from the $WMT_{TA}$.

#### Supplementary Results 3: Hierarchical clustering analysis (HCA)

We next asked whether white matter tracts exhibit higher-order organization based on shared functional profiles. A dendrogram, linking WMTs by their similarity distances, was created as part of the HCA process. The dendrogram's cophenetic correlation coefficient, which measures how faithfully a dendrogram preserves its original pairwise distances, was 0.7446, indicating a strong, high-quality fit. To determine the optimal number of clusters for the dendrogram of WMTs, the Dunn index, an evaluation metric for clustering quality, was computed. It is calculated as the ratio of the minimum inter-cluster distance (separation) to the maximum intra-cluster distance (diameter). A higher Dunn Index value indicates better clustering quality. The analysis favored a five-cluster solution (DI = 0.2972) over other alternatives, suggesting a well-separated and compact cluster structure (**Supplementary Fig. 10**). A list of WMTs within each cluster and their anatomical locations is provided in main manuscript **Fig. 2**.

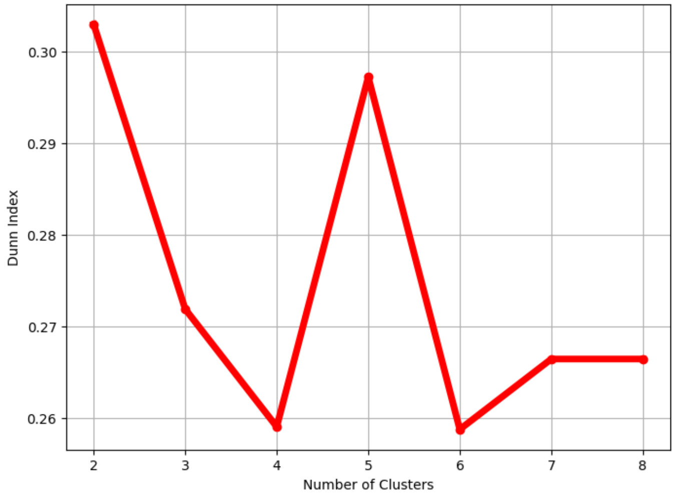


**Supplementary Figure 10. Evaluation of hierarchical clustering solutions.** Quantitative assessment of hierarchical clustering solutions demonstrated that a five-cluster solution provided the optimal balance between cluster compactness and separation, yielding the highest Dunn Index value (DI = 0.2972) relative to the four- and six-cluster alternatives. These results support the selection of the five-cluster hierarchical clustering solution presented in **Fig. 2**.

#### Hierarchical Cluster Detailed Description

To identify shared functional organization among white matter tracts (WMTs) in the termination atlas, tracts were grouped into clusters based on similarity in their functional profiles, such that within-cluster similarity exceeded between-cluster similarity. We then examined the functional profiles of the combined tracts within each cluster and evaluated their correspondence with prior literature. The five cluster descriptions below correspond to those presented in Figure 2 of the main manuscript, with additional tract-level detail provided in **Supplementary Fig. 11**.

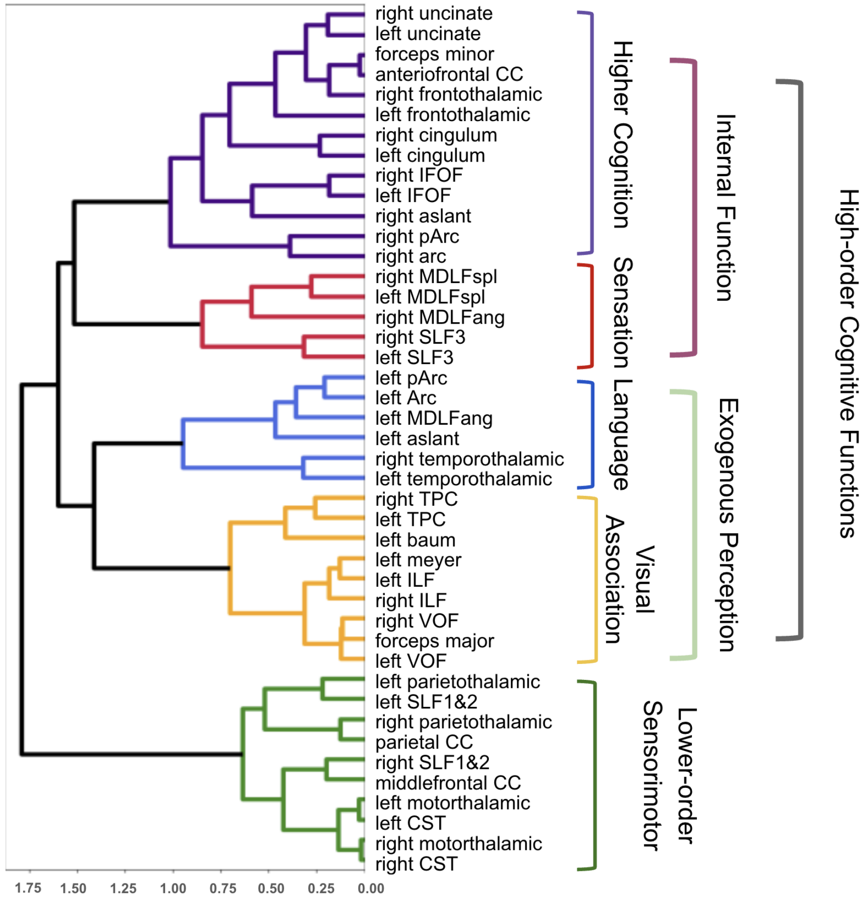


**Supplementary Figure 11. Hierarchical clustering of white matter tracts based on functional associations of tract termination points**. HCA favored a five-cluster solution (Dunn Index = 0.2972), each cluster is visualized with a unique color. The 'height' of a branch splitting along the x-axis indicates the dissimilarity of tract groups, where higher branching corresponds to greater dissimilarity and lower branching indicates less dissimilarity among white matter tracts. Tracts for each cluster are labeled next to their respective branch. Tract Acronyms: CC = corpus callosum, IFOF = Inferior Fronto-Occipital Fasciculus, Arc = arcuate, MDLF = medial longitudinal fasciculus, SLF = Superior longitudinal fasciculus, TPC = Temporo-pontocerebellar, ILF = inferior longitudinal fasciculus, VOF = Vertical Occipital Fasciculus, CST = cortico-spinal tract.

#### Cortical organization and functional characterization of the sensorimotor cluster (Green)

Examining the first cluster (**Fig. 2a** and **Supplementary Fig. 11**, green) revealed that it consisted of 10 tracts: middle frontal CC, parietal CC, and left and right tracts of CST, SLF1 and SLF2, motor thalamic, and parieto-thalamic tracts. The cumulative endpoints for this cluster spread across the dorsal-medial and dorsal-lateral portions of the frontal lobe extending through the supplementary motor, motor cortices, and parietal lobe. There was a high level of symmetry across the right and left hemispheres. Broadly, this cluster was associated with external lower-order cognitive functions, namely motor control. Its functional decoding revealed strong associations with motor functions such as hand movement, movement execution, imagination, and training.

These findings are highly consistent with prior work. For example, early anatomical work examining the CST have identified its as a primary motor pathway that originates from multiple cortical areas, including the primary motor cortex, supplementary motor area, premotor cortex, somatosensory cortex, and cingulate gyrus[27,28]. Additional support for its functional characterization comes from lesion studies showing that damage to the CST can result in motor deficits, highlighting its critical role in voluntary motor control[29]. Similarly, the CC connects homologous areas of the frontal and parietal lobes across hemispheres, which is essential for coordinated bilateral motor activities[30]. Together, functional and anatomical reviews of the SLF highlight the anatomical overlap of this tract with CC and CST along the superior frontal gyrus and the supplementary motor area to the parietal cortices as well as its involvement in supporting the integration of sensory information and motor planning[31].

**Cortical organization and functional characterization of the Visual association cluster (Orange)**

The second cluster (**Fig. 2b** Orange) consisted of 9 tracts: the forceps major, left baum, left meyer, and bilateral tracts of ILF, TPC, and VOF. This cluster, composed of several visual tracts, spreads across the medial and lateral occipital lobe, fusiform gyrus, the inferior temporal gyrus, and the temporal pole (BA 38).

Functional decoding of this cluster revealed its role in extrospective cognitive functions, which are largely related to various aspects of visual processing, including fixation, face perception, gaze control, and saccades. These findings corroborate previous functional understandings of WMTs within this cluster. The VOF has been related to integrating visual information for spatial awareness[7,17,18] and the ILF with object recognition[19–21]. The remaining major visual tracts tend to serve purposes within the occipital lobe. For example, the forceps major supports visual processing by enabling communication between the occipital lobes via the splenium[32]. The baum and meyer's loop are part of the optic radiation and support information transfer to the primary visual cortex[33].

**Cortical organization and functional characterization of the Language cluster (Blue)**

The third cluster (**Fig. 2c** Blue) consisted of 6 tracts: the left and right temporo-thalamic tracts and left tracts of Arcuate, posterior Arcuate, Aslant, and the middle longitudinal fasciculus–superior angular gyrus component (MdLF-Ang)[34]. This cluster was highly left lateralized, with cumulative endpoints of these tracts located in Broca's and Wernicke's area as well as left middle temporal, left inferior temporal gyri, and the left temporal pole. In general, the cluster was associated with higher order cognitive functions, particularly extrospection and language processing. Its functional decoding revealed strong associations with language processing such as semantic processing, sentence comprehension, naming, reading, and speech.

Stroke and lesion research has demonstrated that damage to the left Aslant, one of the WMTs within this cluster, commonly results with speech and language deficits[35] and is more predictive of apraxia than damage to the cortical areas it connects[36]. Further, numerous DTI studies have suggested the role of the MdLF-ang in language processing[37–39]. It is less clear as to why the right temporo-thalamic tracts were a part of this cluster, given their role in auditory processing and memory.

**Cortical organization and functional characterization of the Sensation cluster (Red)**

The fourth cluster (**Fig. 2d** Red) consisted of 5 tracts: bilateral MdLF-spl and SLF3, and the right MdLF-ang. The endpoints of these tracts spread from the ventrolateral frontal cortex, across the sylvian fissure, the temporal-parietal junction, and the sensory cortex. This cluster demonstrated a slight right lateralization and also included portions of the right temporal pole. Its functional decoding was related to processing a wide range of sensory information. The cluster is mainly characterized by associations with processing sensory and proprioceptive information, including tactile (e.g., intensity, touch, pain, heat) and auditory stimuli (sound, auditory, speech). Interoceptive processes such as pain[40,41], balance[42,43], and taste[44] were also noted. Previous studies have also shown a connection between auditory processing and tactile stimuli[45,46].

Recent work has highlighted the importance of major cortico–cortical white matter tracts, particularly the SLF 3 for accurate proprioception after stroke[47], which supports this cluster's relation to touch, movement, and execution. Other work has reported clinical cases supporting the role of MdLF in high-order functions related to acoustic information and further argues that the MdLF may contribute to the learning process associated with verbal-auditory stimuli[48], both of which align with our results linking these tracts to speech, sound, tone, and pitch functions. Direct associations between these tracts and modalities like balance, taste, or responses to acupuncture are less well-established. However, given the involvement of the MdLF and SLF3 in integrating sensory information, they may contribute to the processing of these modalities indirectly.

**Cortical organization and functional characterization of the Higher cognition cluster (Purple)**

The final fifth cluster (**Fig. 2e** Purple) consisted of thirteen WMTs which included bilateral IFOF, bilateral cingulum, bilateral fronto-thalamic tract, bilateral uncinate fasciculus, the anterior frontal CC, forceps minor, right Aslant, right Arcuate, and right posterior Arcuate. Endpoints of these tracts were located medially along the cingulate and frontal lobes, including the mid-frontal gyrus, and demonstrated a resemblance to the default mode network (DMN[49]). This cluster showed a modest right lateralization, which was particularly evident in the right prefrontal cortices and right medial temporal lobe.

Functional decoding of this cluster revealed higher-order cognitive functions related to introspection. More specifically, the cluster was associated with higher-order cognition such as processes related to oneself, moral judgment, decision making, social cognition, and emotion regulation. Many of these functions are commonly associated with DMN[49] and fronto-parietal functions.

Previous studies have demonstrated that the cingulum connects the medial prefrontal cortex (MPFC) and posterior cingulate cortex (PCC), both central nodes of the DMN. This tract is crucial for self-referential thought, emotion regulation, and social cognition. Studies have shown that the cingulum supports the structural integrity necessary for effective DMN connectivity, which underpins these cognitive functions[50,51]. The uncinate fasciculus has been linked to social-emotional processing[52], particularly in patients with the behavioral variant of frontotemporal dementia[53] who demonstrate deficits in social and emotional processing and exhibit uncinate abnormalities[54]. Lastly, the arcuate has been suggested for endogenous attentional control[55].

**Supplementary Results 4: Cluster Convergence with Canonical Functional Networks (Network Correspondence Toolbox)**

We sought to determine how white matter tract cortical terminations would map onto the established canonical functional networks. To this end, we used the white matter tract ensembles and a recently published toolbox that maps 23 different functional network atlases. To determine whether the five tract ensembles identified via HCA correspond to established large-scale cortical brain systems, we quantified their *spatial alignment* with canonical functional networks across multiple atlases[56–63] using the Network Correspondence toolbox (NCT; **Supplementary Fig. 12)**[64].

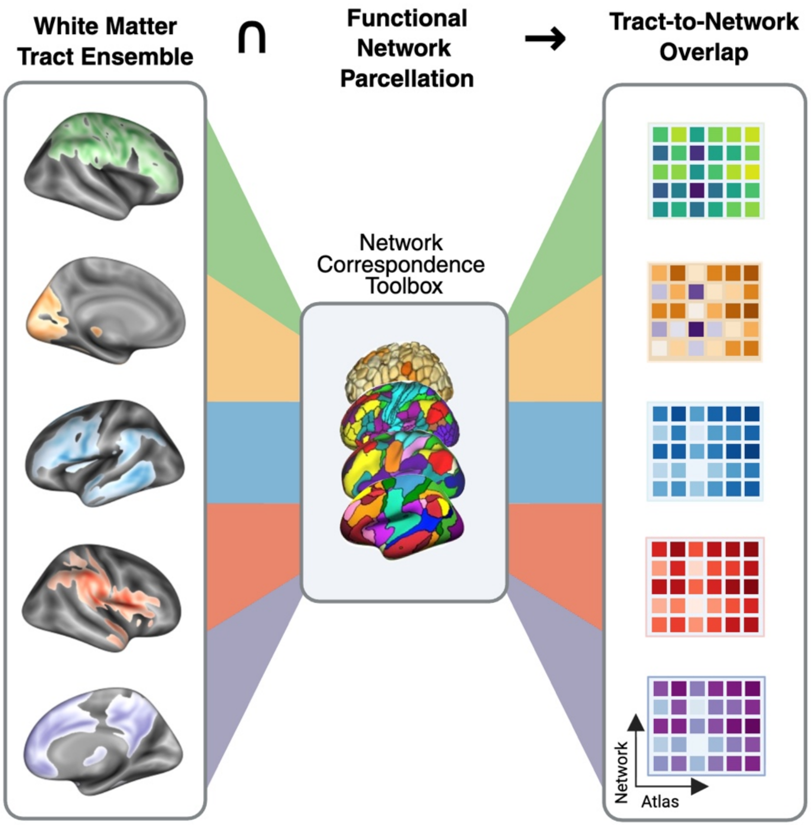


**Supplementary Figure 12. Schematic of network correspondence analysis**. The five white matter tract (WMT) ensembles identified through hierarchical clustering were submitted to the Network Correspondence Toolbox (NCT) to quantify their spatial alignment with canonical cortical functional networks across six widely used brain atlases (EG17[56,57], TY7[58], AS200Yeo17[58,59], MGlasser360J12[60,61], XS268_8[62], and EG286_12[56,63]). For each tract ensemble, the NCT computed spatial correspondence with network parcels using Dice coefficients and evaluated statistical significance using permutation-based spin tests (n = 1,000), generating null distributions against which observed overlap was compared. This framework enabled assessment of whether functionally defined WMT ensembles exhibit non-random correspondence with established large-scale cortical systems. Created in https://BioRender.com

Following the framework described in Kong et al (2025)[64], spatial correspondence between each tract ensemble and each network within a given atlas was quantified, and the statistical significance of this correspondence was evaluated using the NCT. Prior to analysis, all atlas-defined networks and tract-derived cortical termination maps were projected from standard volumetric space to the fsaverage6 surface space to ensure alignment across datasets.

Spatial correspondence was assessed using the Dice coefficient, which ranges from 0 (no overlap) to 1 (complete overlap). To evaluate statistical significance, we employed permutation-based spin tests[65,66] implemented in Python. For each atlas network, 1,000 spatial permutations were generated to construct a null distribution of overlap values, against which observed Dice coefficients were compared **(Supplementary Fig. 13-17; Supplementary Tables 1-6**).

This approach allowed us to assess whether clusters of WMTs, defined by shared functional profiles, systematically map onto known cortical network architectures. Such alignment would indicate that these WMT groupings are not arbitrary, but instead reflect coherent structural organization underlying large-scale cortical systems. Within this framework, overlapping WMTs may support coordinated communication within and between distributed cortical networks, providing a structural basis for their functional integration. The specific networks within each functional atlas that demonstrated significant alignment with the tract ensembles are detailed below.

The ensemble of tracts referred to as "sensorimotor" (**Fig. 2a**, green) exhibited significant correspondence with motor networks, including premotor networks (**Supplementary Fig. 13**; EG17: Premotor dice = 0.1502, p = 0.0340;), somatomotor (EG17: HandSM Premotor dice = 0.1588, p = 0.0450; MG360J12: Somatomotor dice = 0.4870, p = 0.0460; AS200Y17: SomatomotorA dice = 0.2237, p = 0.0380; EG286_12 DorsalSM dice = 0.3022, p = 0.0350), and along with components of the frontoparietal control network (EG17: FrontPar dice = 0.2534, p = 0.0450; AS200Y17: ControlA dice = 0.2237, p = 0.0380, DorsAttnB dice = 0.1831, p = 0.0160; EG286_12 DorsAttn dice = 0.2289, p = 0.0300; TY7 Control dice 0.3208, p= 0.0060). While this ensemble demonstrated a high cortical alignment with the Shen motor network (dice = 0.4962), it did not meet significance. Still, this pattern is consistent with a tract architecture supporting sensorimotor integration and motor planning, aligning with its decoded associations with motor execution and imagery.

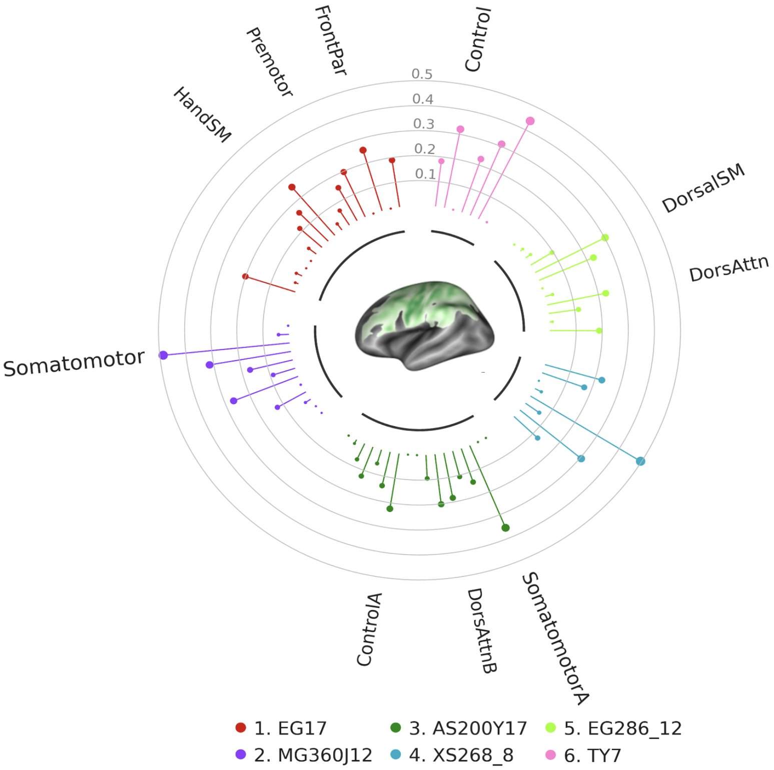


**Supplementary Figure 13. Sensorimotor tract ensemble convergence with Canonical Functional Networks.** The sensorimotor white matter tract cluster ensemble was submitted to the NCT[64] to quantify its cortical correspondence with six common functional brain network atlases[56–63]. Results are visualized using a radial stem plot. The color of the stems corresponds to each of the six functional brain network atlases tested, and the length of the stem corresponds to the dice coefficient between the white matter tract cluster and an individual functional brain network within an atlas. Stems showing a brain network label indicate that the correspondence between the white matter tract cluster and the cortical brain network meets significance (spin tested, 1,000 permutations p<0.05).

The ensemble of tracts referred to as “visual association” (**Fig. 2b**, orange) demonstrated strong overlap with multiple visual networks (**Supplementary Fig. 14**; EG17: LatVis dice = 0.4448, p = 0.0210; MG360J12: Visual1 dice = 0.2231, p =0.0500, Visual2 dice = 0.5528, p = 0.0050; AS200Y17: VisualA dice = 0.3825, p = 0.0240; XS268_8: VisAssoc dice = 0.2241, p = 0.0270, VisualB dice = 0.3787, p = 0.0390; EG286_12 Visual: dice =0.5193 , p = 0.0160; TY7: Visual dice = 0.6044, p =0.0210) and two dorsal attention networks (EG17: DorsAttn dice = 0.2727, p = 0.0420; AS200Y17: DorsAttnA dice = 0.3076, p = 0.0200:), consistent with a posterior pathway system supporting visuospatial processing and attentional orienting[67]. This correspondence aligns with its functional decoding, which emphasizes visual perception, gaze control, and visually guided behavior.

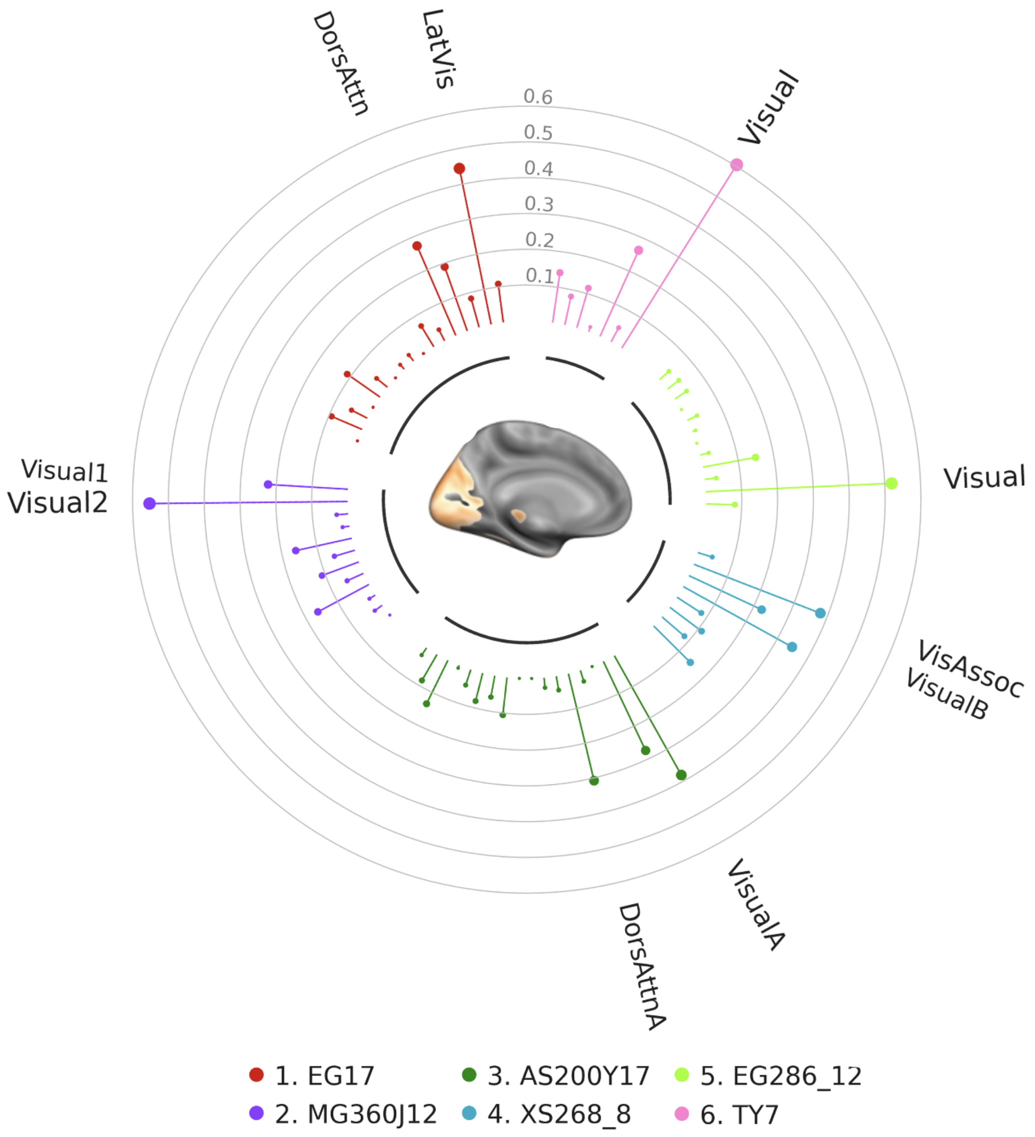


**Supplementary Figure 14. Visual Association tract ensemble convergence with Canonical Functional Networks.** The visual association white matter tract cluster ensemble was submitted to the NCT[64] to quantify its cortical correspondence with six common functional brain network atlases[56–63]. Results are visualized using a radial stem plot where the stem color corresponds to each of the six functional brain network atlases tested and the length of the stem corresponds to the dice coefficient between the white matter tract cluster and an individual functional brain network within an atlas. Stems showing a brain network label indicate that the correspondence between the white matter tract cluster and the cortical brain network meets significance (spin tested, 1,000 permutations p<0.05).

The ensemble of tracts referred to as “language’” (**Fig. 2c**, blue) showed correspondence with a diverse set of frontoparietal (**Supplementary Fig. 15**; EG17: FrontPar dice = 0.2721, p = 0.0040; MG360J12: FrontPar dice = 0.2310, p = 0.0070; XS268 FrontPar dice = 0.3246, p = 0.0010: EG286_12: FrontPar dice = 0.0940, p = 0.0140) and control networks (AS200Y17: ControlA dice = 0.2710, p = 0.0050, ControlB dice = 0.1284, p = 0.0100; TY7: Control dice = 0.2530, p = 0.0030), alongside regions within language (MG360J12: Language dice =0.2190 , p = 0.0090) and default mode networks (AS200Y12: DefaultB dice = 0.2832, p = 0.0360). This distributed pattern is consistent with its decoding profile, highlighting language processing and suggesting a role in integrating linguistic functions with executive and internally directed cognitive processes.

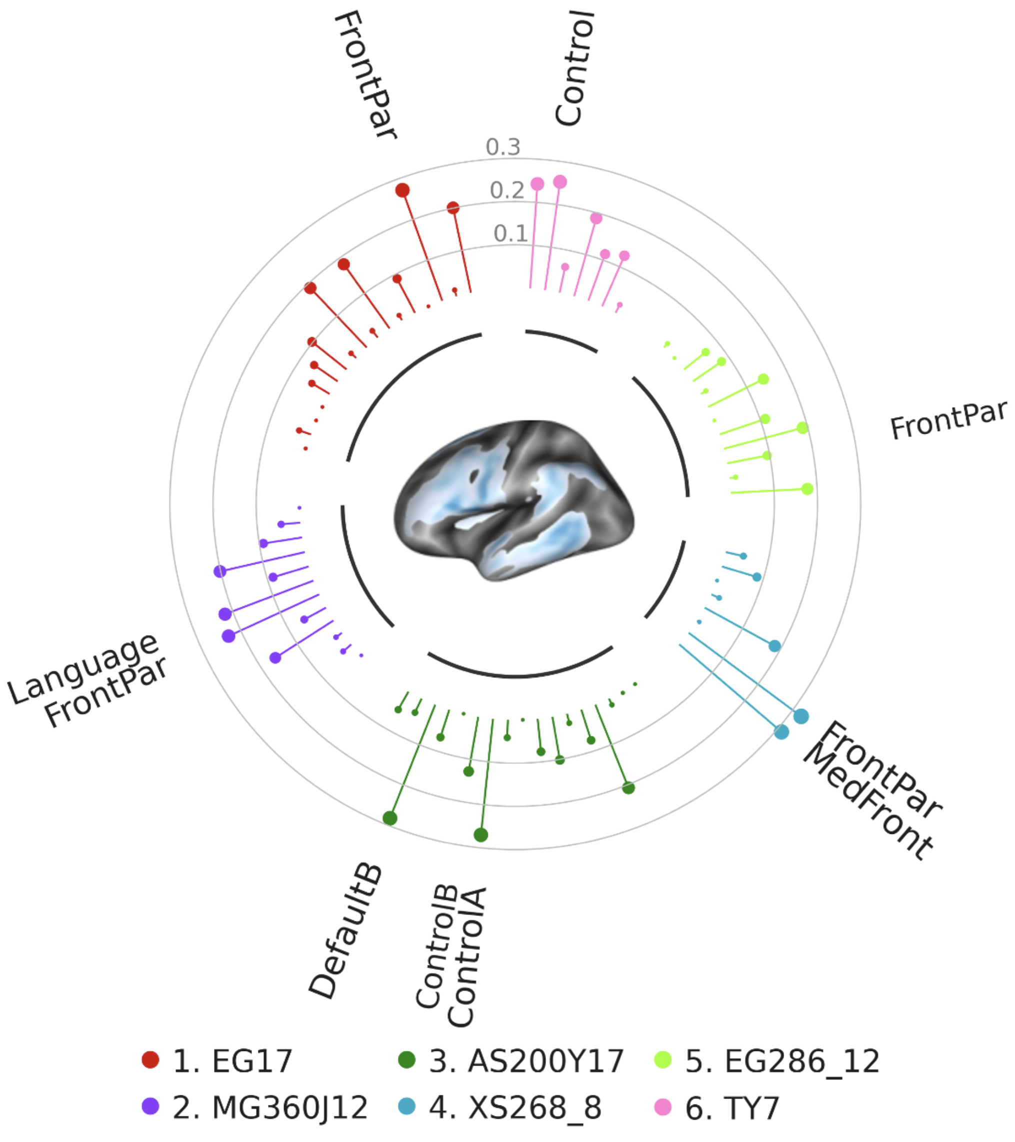


**Supplementary Figure 15. Language tract ensemble convergence with Canonical Functional Networks.** The language white matter tract cluster ensemble was submitted to the NCT[64] to quantify its cortical correspondence with six common functional brain network atlases[56–63]. Results are visualized using a radial stem plot where the stem color corresponds to each of the six functional brain network atlases tested and the length of the stem corresponds to the dice coefficient between the white matter tract cluster and an individual functional brain network within an atlas. Stems showing a brain network label indicate that the correspondence between the white matter tract cluster and the cortical brain network meets significance (spin tested, 1,000 permutations p<0.05).

The ensemble of tracts referred to as “sensation” (**Fig. 2d**, red) showed robust overlap with regions belonging to the cingulo-opercular (**Supplementary Fig. 16**; EG17: CingOperc dice = 0.3416, p = 0.0120; MG360J12: CingOperc dice = 0.3335, p = 0.0320; EG286_12: CingOperc dice = 0.3203, p = 0.0080) and salience/ventral attention networks (AS200Y17: Sal/VenAttnA dice = 0.2428, p = 0.0440; TY17: Sal/VenAttn dice = 0.3229, p = 0.0170), alongside a single motor network (XS268_8: Motor dice = 0.4890, p = 0.0440). This pattern aligns with its decoded associations with sensory, proprioceptive, and interoceptive processes and implicates this cluster in stimulus-driven salience detection and the initiation of contextually appropriate actions[68,69].

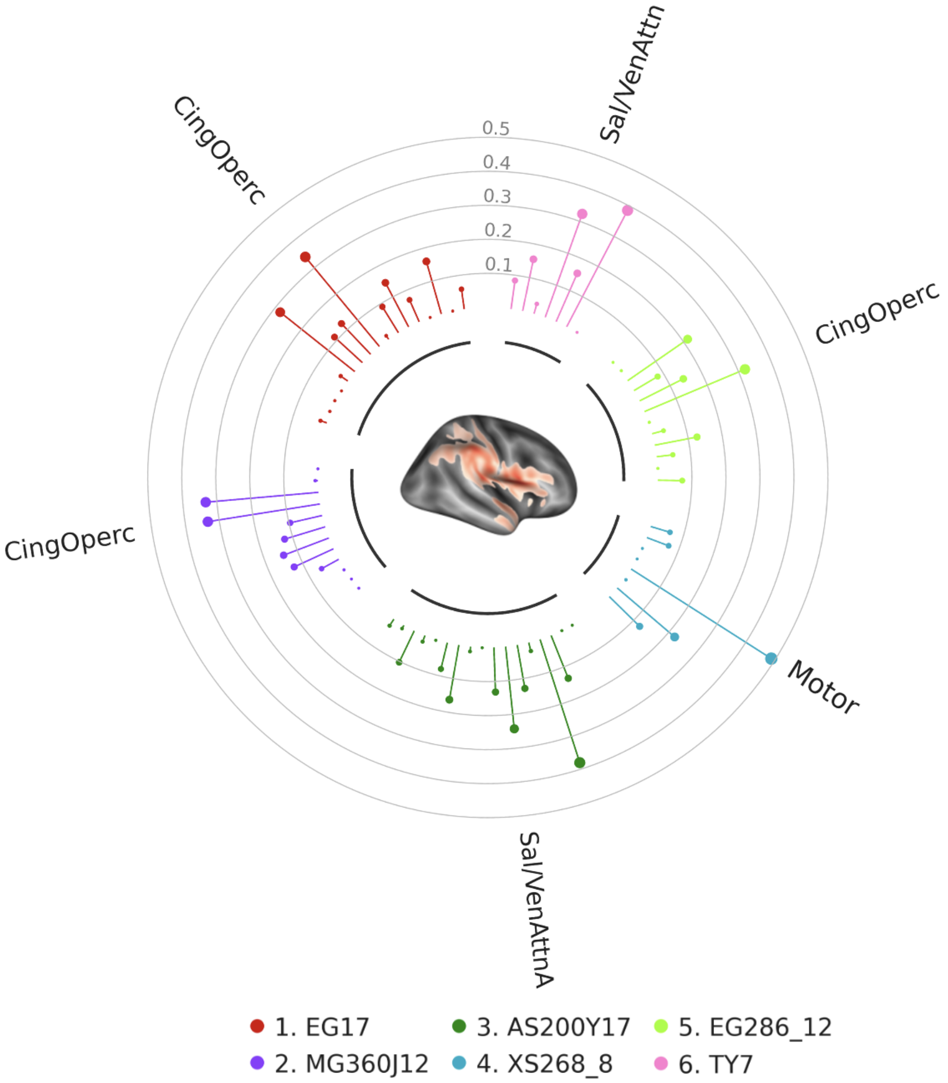


**Supplementary Figure 16. Sensation ensemble convergence with Canonical Functional Networks.** The sensation white matter tract cluster ensemble was submitted to the NCT[64] to quantify its cortical correspondence with six common functional brain network atlases[56–63]. Results are visualized using a radial stem plot where the stem color corresponds to each of the six functional brain network atlases tested and the length of the stem corresponds to the dice coefficient between the white matter tract cluster and an individual functional brain network within an atlas. Stems showing a brain network label indicate that the correspondence between the white matter tract cluster and the cortical brain network meets significance (spin tested, 1,000 permutations p<0.05).

The ensemble of tracts referred to as “higher cognition” (**Fig. 2e**, purple) revealed a notable convergence with the DMN (**Supplementary Fig. 17**; EG17: Default dice = 0.4568, p = 0.0010; MG360J12:Default dice = 0.4269, p = 0.0010; AS200Y17: DefaultA dice = 0.2793, p = 0.0020, DefaultB dice = 0.2441, p = 0.0090; XS268_8: Default dice = 0.2481, p = 0.0130; EG286_12: Default dice = 0.3609, p = 0.0020; TY7: Default dice = 0.4709, p = 0.0020), salience (EG17: Salience dice = 0.1376, p = 0.0020; AS200Y17:Sal/VenAttnB dice = 0.1734, p = 0.0090; XS268_8: SalSubcor dice = 0.3160, p = 0.0020; EG286_12: Salience dice = 0.0141, p = 0.0450), frontal parietal and control (MG360J12: CingOperc dice = 0.3321, p = 0.0230, FrontPar dice = 0.3601, p = 0.0020; AS200Y17:ControlB dice = 0.2457, p = 0.0010; XS268_8: FrontPar dice = 0.3442, p = 0.0320, MedFront dice = 0.2895, p = 0.0110; EG286_12: FrontPar dice = 0.1610, p = 0.0030, TY7: Control dice = 0.3569, p = 0.0040) networks across multiple atlas frameworks. This constellation of associations aligns with the so-called "triple network" model of cognitive control[70], underscoring the integrative role of this cluster in mediating transitions between internal mentation, attentional reorienting, and executive regulation.

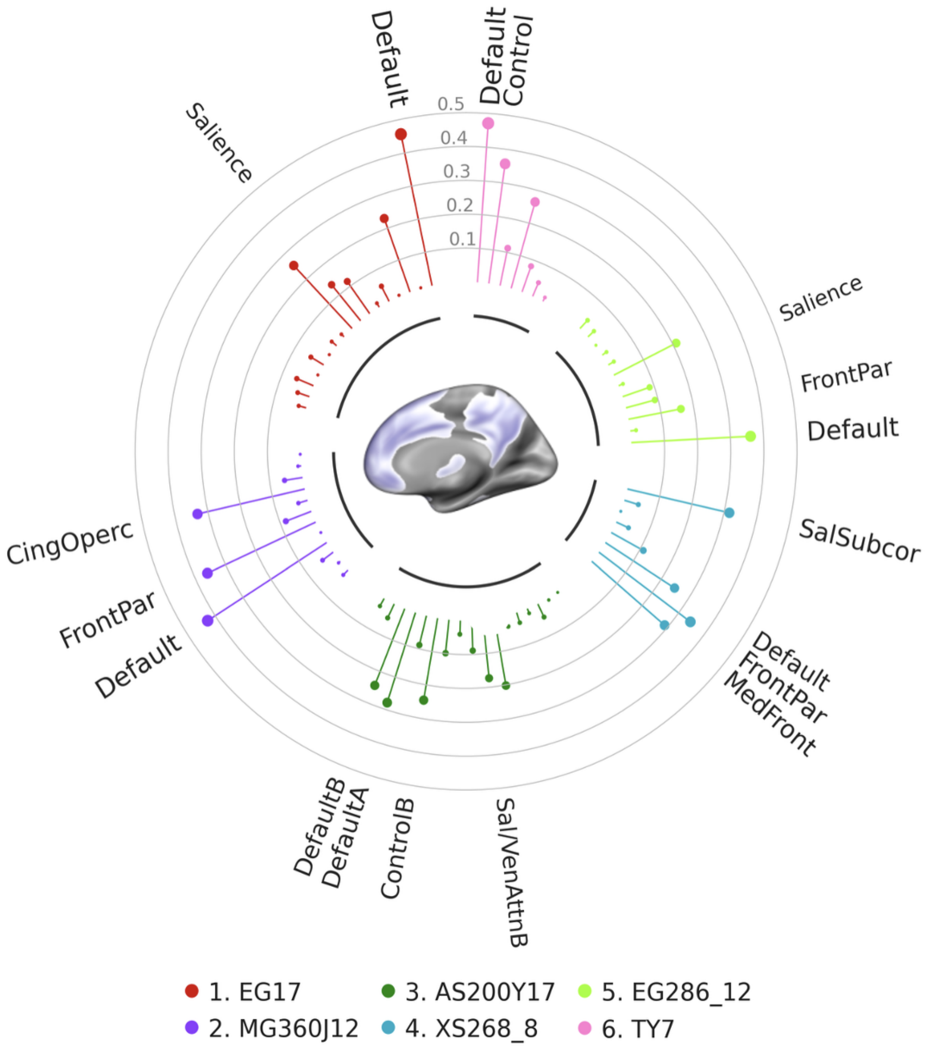


**Supplementary Figure 17. Higher Cognition ensemble convergence with Canonical Functional Networks.** The sensation white matter tract cluster ensemble was submitted to the NCT[64] to quantify its cortical correspondence with six common functional brain network atlases[56–63]. Results are visualized using a radial stem plot where the stem color corresponds to each of the six functional brain network atlases tested and the length of the stem corresponds to the dice coefficient between the white matter tract cluster and an individual functional brain network within an atlas. Stems showing a brain network label indicate that the correspondence between the white matter tract cluster and the cortical brain network meets significance (spin tested, 1,000 permutations p<0.05).

| EG17[57] Network Name | Cluster 1: Sensorimotor | | Cluster 2: Visual Association | | Cluster 3: Language | | Cluster 4: Sensation | | Cluster 5: Higher Cognition | |
|---|---|---|---|---|---|---|---|---|---|---|
| | dice | p_value | dice | p_value | dice | p_value | dice | p_value | dice | p_value |
| Default | 0.1896 | 0.6274 | 0.1083 | 0.8372 | 0.2004 | 0.2118 | 0.0589 | 0.9481 | 0.4568 | **0.0010** |
| LatVis | 0.0000 | 0.9700 | 0.4448 | **0.0210** | 0.0149 | 0.7962 | 0.0000 | 0.8581 | 0.0017 | 0.9980 |
| FrontPar | 0.2534 | **0.0450** | 0.0835 | 0.6523 | 0.2721 | **0.0040** | 0.1603 | 0.3067 | 0.2303 | 0.1089 |
| MedVis | 0.0000 | 0.6993 | 0.1900 | 0.0949 | 0.0000 | 0.7512 | 0.0000 | 0.5564 | 0.0032 | 0.8601 |
| DorsAttn | 0.1975 | 0.2258 | 0.2727 | **0.0420** | 0.0891 | 0.5055 | 0.0705 | 0.5245 | 0.0502 | 0.9131 |
| Premotor | 0.1502 | **0.0340** | 0.0346 | 0.4875 | 0.0130 | 0.7103 | 0.1468 | 0.1229 | 0.0135 | 0.9401 |
| Language | 0.0688 | 0.6474 | 0.0679 | 0.5225 | 0.1823 | 0.0659 | 0.0901 | 0.4046 | 0.1164 | 0.3467 |
| Salience | 0.0302 | 0.6384 | 0.0000 | 0.7582 | 0.0200 | 0.6903 | 0.0125 | 0.5804 | 0.1376 | **0.0020** |
| CingOperc | 0.2537 | 0.2038 | 0.0215 | 0.9351 | 0.1896 | 0.1119 | 0.3416 | **0.0120** | 0.2561 | 0.1049 |
| HandSM | 0.1588 | **0.0450** | 0.0169 | 0.5854 | 0.0162 | 0.7073 | 0.1242 | 0.1778 | 0.0109 | 0.9281 |
| FaceSM | 0.1140 | 0.2048 | 0.0000 | 0.5834 | 0.1020 | 0.1558 | 0.1111 | 0.1608 | 0.0192 | 0.6184 |
| Auditory | 0.0366 | 0.5654 | 0.0390 | 0.4016 | 0.0662 | 0.4555 | 0.2801 | 0.0869 | 0.0019 | 0.9660 |
| AntMTL | 0.0000 | 0.7463 | 0.1118 | 0.2168 | 0.0478 | 0.4476 | 0.0253 | 0.4056 | 0.0450 | 0.4855 |
| PostMTL | 0.0000 | 0.7223 | 0.0005 | 0.4645 | 0.0000 | 0.6643 | 0.0000 | 0.5644 | 0.0010 | 0.8132 |
| ParMemory | 0.0243 | 0.5015 | 0.0494 | 0.2907 | 0.0000 | 0.7572 | 0.0034 | 0.5215 | 0.0544 | 0.1648 |
| Context | 0.0127 | 0.7742 | 0.0922 | 0.1608 | 0.0288 | 0.6533 | 0.0035 | 0.7962 | 0.0371 | 0.5165 |
| FootSM | 0.2010 | 0.0679 | 0.0002 | 0.6164 | 0.0023 | 0.6903 | 0.0197 | 0.4396 | 0.0230 | 0.7293 |

**Supplementary Table 1: White matter tract Cluster spatial alignment with Gordon canonical functional networks.** White matter tract endpoint clusters from the HCA five-cluster solution were submitted to the NCT [64] to quantify their cortical correspondence with the EG17[57] functional atlas. Significant findings are emphasized with bold text.

| MG360J12[60,61] Network Name | Cluster 1: Sensorimotor | | Cluster 2: Visual Association | | Cluster 3: Language | | Cluster 4: Sensation | | Cluster 5: Higher Cognition | |
|---|---|---|---|---|---|---|---|---|---|---|
| | dice | p_value | dice | p_value | dice | p_value | dice | p_value | dice | p_value |
| Visual1 | 0.0036 | 0.6923 | 0.2231 | **0.0500** | 0.0000 | 0.8222 | 0.0004 | 0.6144 | 0.0011 | 0.9560 |
| Visual2 | 0.0403 | 0.7093 | 0.5528 | **0.0050** | 0.0443 | 0.7413 | 0.0077 | 0.7203 | 0.0095 | 0.9980 |
| Somatomotor | 0.4870 | **0.0460** | 0.0322 | 0.7532 | 0.0906 | 0.6424 | 0.3328 | 0.1778 | 0.0549 | 0.9720 |
| CingOperc | 0.3160 | 0.1449 | 0.0197 | 0.9401 | 0.2015 | 0.0979 | 0.3335 | **0.0320** | 0.3321 | **0.0230** |
| DorsAttn | 0.1681 | 0.1808 | 0.1602 | 0.1698 | 0.0859 | 0.4436 | 0.0964 | 0.3776 | 0.0290 | 0.9790 |
| Language | 0.0882 | 0.4825 | 0.0596 | 0.5514 | 0.2190 | **0.0090** | 0.1244 | 0.2378 | 0.0821 | 0.4286 |
| FrontPar | 0.2660 | 0.0949 | 0.1097 | 0.7283 | 0.2310 | **0.0070** | 0.1425 | 0.4855 | 0.3601 | **0.0020** |
| Auditory | 0.0045 | 0.6334 | 0.0503 | 0.3037 | 0.0572 | 0.3996 | 0.1273 | 0.1508 | 0.0000 | 0.9111 |
| Default | 0.1256 | 0.9321 | 0.1620 | 0.6523 | 0.1600 | 0.5325 | 0.0575 | 0.9371 | 0.4269 | **0.0010** |
| PostMulti | 0.0238 | 0.5544 | 0.0190 | 0.4855 | 0.0172 | 0.5225 | 0.0026 | 0.6563 | 0.0398 | 0.2937 |
| VentMulti | 0.0000 | 0.6703 | 0.0254 | 0.3427 | 0.0247 | 0.4555 | 0.0000 | 0.5305 | 0.0070 | 0.7203 |
| OrbitAffective | 0.0000 | 0.6723 | 0.0000 | 0.4286 | 0.0000 | 0.6643 | 0.0000 | 0.5095 | 0.0201 | 0.4116 |

**Supplementary Table 2: White matter tract Cluster spatial alignment with Glasser canonical functional networks.** White matter tract endpoint clusters from the HCA five-cluster solution were submitted to the NCT [64] to quantify their cortical correspondence with the MG360J12[60,61] functional atlas. Significant findings are emphasized with bold text.

| AS200Y17 Network Name | Cluster 1: Sensorimotor | | Cluster 2: Visual Association | | Cluster 3: Language | | Cluster 4: Sensation | | Cluster 5: Higher Cognition | |
|---|---|---|---|---|---|---|---|---|---|---|
| | dice | p_value | dice | p_value | dice | p_value | dice | p_value | dice | p_value |
| TempPar | 0.0015 | 0.7123 | 0.0240 | 0.4056 | 0.0478 | 0.3976 | 0.0227 | 0.4156 | 0.0256 | 0.6603 |
| DefaultC | 0.0173 | 0.7622 | 0.0838 | 0.2438 | 0.0358 | 0.6214 | 0.0107 | 0.6484 | 0.0457 | 0.4975 |
| DefaultB | 0.0696 | 0.7642 | 0.1352 | 0.4096 | 0.2832 | **0.0360** | 0.1026 | 0.4605 | 0.2441 | **0.0090** |
| DefaultA | 0.1250 | 0.4206 | 0.0081 | 0.9291 | 0.0670 | 0.7612 | 0.0192 | 0.8721 | 0.2793 | **0.0020** |
| ControlC | 0.0577 | 0.3397 | 0.0454 | 0.3347 | 0.0000 | 0.7842 | 0.0024 | 0.5514 | 0.0889 | 0.1149 |
| ControlB | 0.1385 | 0.2258 | 0.0808 | 0.5574 | 0.1284 | **0.0100** | 0.0795 | 0.5245 | 0.2457 | **0.0010** |
| ControlA | 0.2237 | **0.0380** | 0.0607 | 0.6284 | 0.2701 | **0.0050** | 0.1629 | 0.1968 | 0.0992 | 0.4695 |
| LimbicA | 0.0000 | 0.7612 | 0.1053 | 0.2607 | 0.0411 | 0.5035 | 0.0142 | 0.4835 | 0.0402 | 0.5834 |
| LimbicB | 0.0002 | 0.7283 | 0.0000 | 0.5295 | 0.0000 | 0.6653 | 0.0007 | 0.5504 | 0.0887 | 0.2378 |
| Sal/VenAttnB | 0.0942 | 0.3816 | 0.0000 | 0.8921 | 0.0765 | 0.2627 | 0.1311 | 0.1858 | 0.1734 | **0.0090** |
| Sal/VenAttnA | 0.2018 | 0.0859 | 0.0279 | 0.8931 | 0.1014 | 0.3327 | 0.2428 | **0.0440** | 0.2020 | 0.1419 |
| DorsAttnB | 0.1831 | **0.0160** | 0.0397 | 0.5405 | 0.0223 | 0.6883 | 0.1295 | 0.2108 | 0.0323 | 0.7912 |
| DorsAttnA | 0.1065 | 0.4925 | 0.3076 | **0.0200** | 0.0744 | 0.5245 | 0.0256 | 0.5644 | 0.0313 | 0.8561 |
| SomatomotorB | 0.1417 | 0.4326 | 0.0321 | 0.5445 | 0.2073 | 0.1708 | 0.3808 | 0.0639 | 0.0147 | 0.9471 |
| SomatomotorA | 0.3571 | **0.0350** | 0.0007 | 0.7383 | 0.0160 | 0.7622 | 0.1359 | 0.3137 | 0.0430 | 0.8651 |
| VisualB | 0.0005 | 0.7652 | 0.2756 | 0.0729 | 0.0031 | 0.8222 | 0.0000 | 0.6613 | 0.0050 | 0.9421 |
| VisualA | 0.0006 | 0.7682 | 0.3825 | **0.0240** | 0.0008 | 0.8042 | 0.0003 | 0.6573 | 0.0000 | 0.9910 |

**Supplementary Table 3: White matter tract Cluster spatial alignment with Schaefer canonical functional networks.** White matter tract endpoint clusters from the HCA five-cluster solution were submitted to the NCT [64] to quantify their cortical correspondence with the AS200Y17[58,59] functional atlas. Significant findings are emphasized with bold text.

| XS268_8[62] Network Name | Cluster 1: Sensorimotor | | Cluster 2: Visual Association | | Cluster 3: Language | | Cluster 4: Sensation | | Cluster 5: Higher Cognition | |
|---|---|---|---|---|---|---|---|---|---|---|
| | dice | p_value | dice | p_value | dice | p_value | dice | p_value | dice | p_value |
| MedFront | 0.1252 | 0.7912 | 0.1445 | 0.5694 | 0.3114 | **0.0210** | 0.1252 | 0.5754 | 0.2895 | **0.0110** |
| FrontPar | 0.3050 | 0.1818 | 0.0822 | 0.7602 | 0.3246 | **0.0010** | 0.2221 | 0.2677 | 0.3442 | **0.0320** |
| Default | 0.0650 | 0.8811 | 0.1109 | 0.5195 | 0.0062 | 0.9630 | 0.0053 | 0.9800 | 0.2481 | **0.0130** |
| Motor | 0.4962 | 0.1828 | 0.0810 | 0.8352 | 0.1849 | 0.4795 | 0.4890 | **0.0440** | 0.1104 | 0.9790 |
| VisualA | 0.0277 | 0.6983 | 0.3473 | 0.0769 | 0.0198 | 0.8022 | 0.0000 | 0.7692 | 0.0402 | 0.8422 |
| VisualB | 0.0000 | 0.6903 | 0.2241 | **0.0270** | 0.0000 | 0.7203 | 0.0000 | 0.5335 | 0.0000 | 0.9201 |
| VisAssoc | 0.1711 | 0.4256 | 0.3787 | **0.0390** | 0.0846 | 0.5684 | 0.0682 | 0.5235 | 0.0438 | 0.8981 |
| SalSubcor | 0.2265 | 0.3397 | 0.0419 | 0.90 | 0.0418 | 0.9520 | 0.0592 | 0.8242 | 0.3160 | **0.0020** |

**Supplementary Table 4: White matter tract Cluster spatial alignment with Shen canonical functional networks.** White matter tract endpoint clusters from the HCA five-cluster solution were submitted to the NCT [64] to quantify their cortical correspondence with the functional atlas. Significant findings are emphasized with bold text.

| EG286_12[63] Network Name | Cluster 1: Sensorimotor | | Cluster 2: Visual Association | | Cluster 3: Language | | Cluster 4: Sensation | | Cluster 5: Higher Cognition | |
|---|---|---|---|---|---|---|---|---|---|---|
| | dice | p_value | dice | p_value | dice | p_value | dice | p_value | dice | p_value |
| Default | 0.1885 | 0.3307 | 0.0809 | 0.81 | 0.1774 | 0.2058 | 0.0717 | 0.8132 | 0.3609 | **0.0020** |
| Visual | 0.0106 | 0.8192 | 0.5193 | **0.0160** | 0.0140 | 0.8322 | 0.0006 | 0.8222 | 0.0175 | 0.9970 |
| FrontPar | 0.1148 | 0.1009 | 0.0318 | 0.6284 | 0.0940 | **0.0140** | 0.0480 | 0.4605 | 0.1610 | **0.0030** |
| DorsAttn | 0.2289 | **0.0300** | 0.1493 | 0.2877 | 0.1887 | 0.0509 | 0.1264 | 0.3526 | 0.0901 | 0.6444 |
| VentAttn | 0.0292 | 0.7113 | 0.0243 | 0.6124 | 0.1114 | 0.2078 | 0.0345 | 0.5225 | 0.0857 | 0.2368 |
| Salience | 0.0000 | 0.7473 | 0.0000 | 0.5375 | 0.0000 | 0.6454 | 0.0015 | 0.4865 | 0.0141 | **0.0450** |
| CingOperc | 0.2274 | 0.1558 | 0.0093 | 0.9461 | 0.1431 | 0.2038 | 0.3203 | **0.0080** | 0.2104 | 0.1558 |
| DorsalSM | 0.3022 | **0.0350** | 0.0297 | 0.5774 | 0.0132 | 0.7662 | 0.1437 | 0.2727 | 0.0190 | 0.9750 |
| VentralSM | 0.0957 | 0.0529 | 0.0000 | 0.4875 | 0.0805 | 0.1768 | 0.0808 | 0.1808 | 0.0157 | 0.5894 |
| Auditory | 0.0226 | 0.5864 | 0.0396 | 0.3756 | 0.0638 | 0.3846 | 0.2143 | 0.0749 | 0.0019 | 0.9670 |
| MedPar | 0.0126 | 0.5065 | 0.0398 | 0.1678 | 0.0000 | 0.7293 | 0.0026 | 0.4825 | 0.0245 | 0.2488 |
| ParOcc | 0.0000 | 0.7273 | 0.0347 | 0.3397 | 0.0121 | 0.6234 | 0.0000 | 0.6194 | 0.0332 | 0.3157 |

**Supplementary Table 5: White matter tract Cluster spatial alignment with Gordon canonical functional networks.** White matter tract endpoint clusters from the HCA five-cluster solution were submitted to the NCT [64] to quantify their cortical correspondence with the EG286_12[63] functional atlas. Significant findings are emphasized with bold text.

| TY7[58] Network Name | Cluster 1: Sensorimotor | | Cluster 2: Visual Association | | Cluster 3: Language | | Cluster 4: Sensation | | Cluster 5: Higher Cognition | |
|---|---|---|---|---|---|---|---|---|---|---|
| | dice | p_value | dice | p_value | dice | p_value | dice | p_value | dice | p_value |
| Visual | 0.0043 | 0.8312 | 0.6044 | **0.0210** | 0.0208 | 0.8362 | 0.0008 | 0.7942 | 0.0129 | 0.9990 |
| Somatomotor | 0.4398 | 0.1309 | 0.0448 | 0.7892 | 0.1279 | 0.5954 | 0.3861 | 0.1049 | 0.0436 | 0.9900 |
| DorsAttn | 0.3093 | 0.1179 | 0.2639 | 0.1628 | 0.1147 | 0.4905 | 0.1552 | 0.3497 | 0.0804 | 0.8701 |
| Sal/VenAttn | 0.2253 | 0.2777 | 0.0143 | 0.9630 | 0.1878 | 0.0929 | 0.3229 | **0.0170** | 0.2652 | 0.0719 |
| Limbic | 0.0000 | 0.8731 | 0.1158 | 0.3706 | 0.0608 | 0.6084 | 0.0300 | 0.5465 | 0.1125 | 0.3796 |
| Control | 0.3208 | **0.0060** | 0.0814 | 0.7992 | 0.2530 | **0.0030** | 0.1552 | 0.3706 | 0.3569 | **0.0040** |
| Default | 0.1827 | 0.9201 | 0.1412 | 0.7982 | 0.2427 | 0.1758 | 0.0843 | 0.9161 | 0.4709 | **0.0020** |

**Supplementary Table 6: White matter tract Cluster spatial alignment with Yeo canonical functional networks.** White matter tract endpoint clusters from the HCA five-cluster solution were submitted to the NCT [64] to quantify their cortical correspondence with the TY7[58] functional atlas. Significant findings are emphasized with bold text.

**Supplementary Methods**

Understanding the cortical termination points of white matter tracts is essential for elucidating the structural organization of the human brain and its relationship to function. While existing white matter atlases provide valuable insights into major fiber pathways, they generally lack systematic mappings of where these tracts interface with the cortical surface. To address this gap, we generated a white matter tract termination atlas (WMTTA) by projecting tractography-derived streamline endpoints onto the cortical surface and aggregating these maps across subjects. This approach enables a more precise characterization of structure–function relationships by linking white matter pathways to their cortical targets. To ensure reproducibility and scalability, all processing steps were implemented using standardized applications on brainlife.io. **Supplementary Table 7** provides a complete list of the brainlife.io applications and version numbers used to generate the $WMT_{TA}$.

| Name | Version | Brainlife DOI |
| --- | --- | --- |
| Remove tract outliers | 1.4 | 10.25663/brainlife.app.195 |
| Generate endpoint map | 1.0 | 10.25663/brainlife.app.194 |
| Reslice ROIs to match input anatomy | 1.0 | 10.25663/brainlife.app.522 |
| Compute and warp from subject-to-standard space | 1.0 | 10.25663/brainlife.app.670 |
| Map termination ROIs to cortical surface | 1.0 | 10.25663/brainlife.app.759 |
| Smooth data mapped to cortical surface | 1.0 | 10.25663/brainlife.app.768 |
| Compute average cortexmap data for group analysis | 1.1 | 10.25663/brainlife.app.552 |

**Supplementary Table 7. brainlife.io apps used to generate the $WMT_{TA}$.**
